# Predicting *Vibrio cholerae* infection and disease severity using metagenomics in a prospective cohort study

**DOI:** 10.1101/2020.02.25.960930

**Authors:** Inès Levade, Morteza M. Saber, Firas Midani, Fahima Chowdhury, Ashraful I. Khan, Yasmin A. Begum, Edward T. Ryan, Lawrence A. David, Stephen B. Calderwood, Jason B. Harris, Regina C. LaRocque, Firdausi Qadri, B. Jesse Shapiro, Ana A. Weil

**Affiliations:** Department of Biological Sciences, University of Montreal, Montreal, QC, Canada; Program in Computational Biology and Bioinformatics, Duke University, Durham, NC, USA; Center for Genomic and Computational Biology, Duke University, Durham, NC, USA; Department of Molecular Genetics and Microbiology, Duke University, Durham, NC, USA; Center for Vaccine Sciences, International Centre for Diarrhoeal Disease Research, Dhaka, Bangladesh; Division of Infectious Diseases, Massachusetts General Hospital, Boston, MA, USA; Department of Medicine, Harvard Medical School, Boston, MA USA; Department of Biomedical Engineering, Duke University, Durham, NC, USA; Department of Immunology and Infectious Diseases, Harvard T.H. Chan School of Public Health, Boston, MA, USA; Department of Microbiology, Harvard Medical School, Boston, MA USA; Department of Pediatrics, Harvard Medical School, Boston, MA, USA; Department of Microbiology and Immunology, McGill University, Montreal, QC, Canada; McGill Genome Centre, Montreal, QC, Canada; Division of Allergy and Infectious Diseases, University of Washington, Seattle, WA, USA

**Keywords:** *Vibrio cholerae*, cholera, microbiome, machine learning, metagenomics

## Abstract

**Background:** Susceptibility to *Vibrio cholerae* infection is impacted by blood group, age, and pre-existing immunity, but these factors only partially explain who becomes infected. A recent study used 16S rRNA amplicon sequencing to quantify the composition of the gut microbiome and identify predictive biomarkers of infection with limited taxonomic resolution.

**Methods:** To achieve increased resolution of gut microbial factors associated with *V. cholerae* susceptibility and identify predictors of symptomatic disease, we applied deep shotgun metagenomic sequencing to a cohort of household contacts of patients with cholera.

**Results:** Using machine learning, we resolved species, strains, gene families, and cellular pathways in the microbiome at the time of exposure to *V. cholerae* to identify markers that predict infection and symptoms. Use of metagenomic features improved the precision and accuracy of prediction relative to 16S sequencing. We also predicted disease severity, although with greater uncertainty than our infection prediction. Species within the genera *Prevotella* and *Bifidobacterium* predicted protection from infection, and genes involved in iron metabolism also correlated with protection.

**Conclusion:** Our results highlight the power of metagenomics to predict disease outcomes and suggest specific species and genes for experimental testing to investigate mechanisms of microbiome-related protection from cholera.

**SUMMARY:** Cholera infection and disease severity can be predicted using metagenomic sequencing of the gut microbiome pre-infection in a prospective cohort, and suggests potentially protective bacterial species and genes.

## INTRODUCTION

Cholera is an acute diarrheal disease caused by *Vibrio cholerae*. It is a major public health threat worldwide that continues to cause major outbreaks, such as in Yemen, where over 1.7 million cases have been reported since 2016 (1,2). Transmission of *V. cholerae* between household members commonly occurs through shared sources of contaminated food or water or through fecal-oral spread (3,4). The clinical spectrum of disease ranges from asymptomatic infection to severe watery diarrhea that can lead to fatal dehydration (5). Host factors such as age, innate immune factors, blood group, or prior acquired immunity partially explain why some people are more susceptible to *V. cholerae* infection than others, but a substantial amount of the variation remains unexplained (6).

The gut bacterial community can protect against enteropathogenic infections (7), and may explain some of the variation in *V. cholerae* susceptibility. Several studies have identified commensal bacteria and mechanisms that could be protective against *V. cholerae*. For instance, a species enriched in the gut microbiota of patients recovering from cholera, *Blautia obeum*, was found to interfere with *V. cholerae* pathogenicity through quorum-sensing inhibition in a mouse model (8). Other experiments have demonstrated that alteration of commensal-derived metabolite levels influenced host susceptibility by affecting *V. cholerae* growth or colonization (9-13).

Studies of *V. cholerae* and the gut microbiota often focus on a few bacterial species, or involve patients who already have symptomatic cholera (8,14). One study recently characterized the gut microbiome of healthy individuals exposed to *V. cholerae* (15). In this study, Midani *et al*. developed a machine learning model to predict susceptibility based on 16S rRNA gene amplicon sequencing of the gut microbiota in a group known to have high risk of infection: household contacts of confirmed cholera patients (4). Midani *et al* showed that microbiome composition at the time *V. cholerae* exposure to can predict infection with similar or better accuracy as commonly measured host factors known to impact susceptibility. However, 16S rRNA sequencing has limited taxonomic resolution and does not identify the genetic mechanisms of protection.

Here we used shotgun metagenomics to analyze an expanded prospective cohort of persons exposed to *V. cholerae* in Bangladesh. Our metagenomic analysis yielded improved outcome predictions compared to 16S rRNA sequencing, and identified bacterial genes associated with remaining uninfected after exposure to *V. cholerae*. We are also able to predict disease severity among infected contacts, albeit with lower power and precision than susceptibility. Finally, we highlight several microbiome-encoded metabolic functions associated with protection against cholera.

## METHODS

### Sample collection, clinical outcomes and metagenomic sequencing

As described in (15), household contacts were enrolled within 6 hours of the presentation of an index cholera case at the icddr,b (International Center for Diarrheal Disease Research, Bangladesh) Dhaka Hospital. Index patients with severe acute diarrhea, a stool culture positive for *V. cholerae*, age between 2 and 60 years old, and no major comorbid conditions were recruited (4,6). A clinical assessment of symptoms in household contacts was conducted daily for the 10-day period after presentation of the index case, and repeated on day 30. We collected demographic information, rectal swabs, and blood samples for ABO typing and vibriocidal antibody titers as described in the Supplementary Methods. During the observation period, contacts were determined to be infected if any rectal swab culture was positive for *V. cholerae* and/or if the contact developed diarrhea and a 4-fold increase in vibriocidal titer during the follow-up period (4,6). Contacts with positive rectal swabs developing watery diarrhea were categorized as symptomatic and those without diarrhea were considered asymptomatic (**Figure 1**). *V. cholerae* positive contacts (by culture or deep 16S amplicon sequencing (15)) at the time of enrollment were excluded, in addition to contacts who reported antibiotic use or diarrhea during the week prior to enrollment. DNA extraction was performed for the selected samples and used for shotgun metagenomics sequencing. Details on cohorts, sequencing methods and sample processing are described in Supplementary Methods.

**Figure 1.**
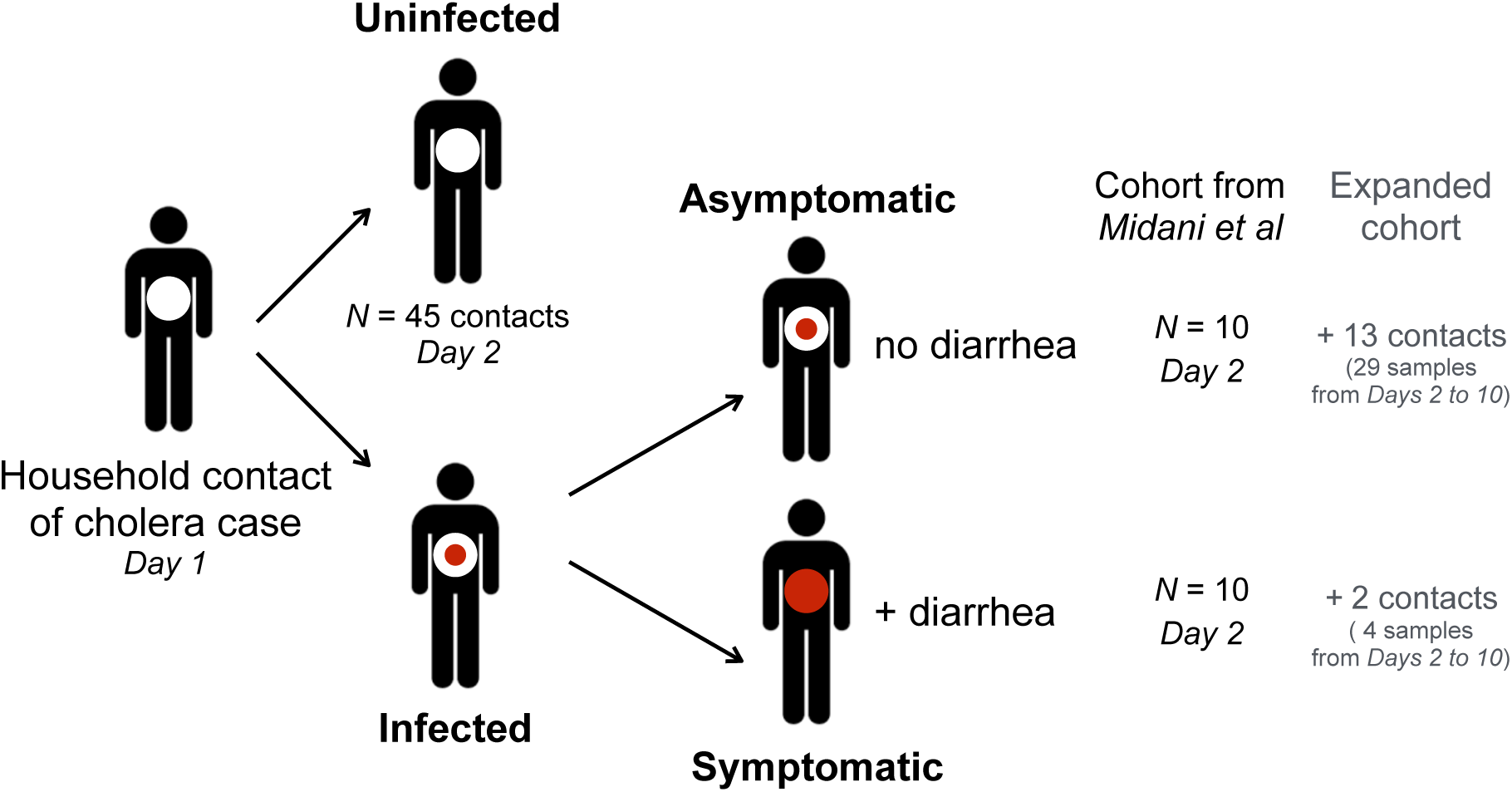
Study cohort in Dhaka, Bangladesh. After presentation of a *V. cholerae* culture-positive index case to the hospital on day 1, household contacts were enrolled on day 2. The expanded cohort includes the Midani 2018 cohort (15), with an addition of 33 samples from infected individuals (13 asymptomatic and 2 symptomatic).

### Taxonomic/functional profiling and predictive model construction

We used MetaPhlAn2 (version 2.9) (16) for taxonomic profiling and HUMAnN2 (17) to profile cellular pathways (from MetaCyc) and gene families (identified using the PFAM database). For identification of biomarkers of susceptibility and disease severity, we used MetAML (18) to apply a random forests (RF) classifier on species, pathways and gene-family relative abundances, as well as strain-specific markers presence/absence. Models constructed using each of these features types were compared to a random dataset with shuffled labels, and to a model constructed with clinical/demographic data, using two-sample, two-sided *t*-tests over 20 replicate cross-validation (18). We used a stratified 3-fold cross validation approach, splitting our dataset into validation and training sets (1/3 and 2/3 of samples, respectively) with the same infected:uninfected ratio. We used an embedded feature selection strategy to identify the most useful features and improve model accuracy. Feature relative importance was computed using the mean decrease in impurity strategy, which calculates importance of each feature as the sum of the number of nodes (across all trees) that use the feature, proportional to the number of samples each of these nodes splits (18). Further details are described in the Supplementary Methods.

### Data availability

After removal of human reads (Supplementary Methods), the sequence data has been deposited in NCBI under BioProject PRJNA608678.

## RESULTS

### Metagenomic sequencing of the gut microbiome in household contacts exposed to *V. cholerae*

We performed metagenomic sequencing of the gut microbiome in 65 contacts of cholera cases from a cohort described by Midani *et al*. (15), from which sufficient DNA remained. Of these 65 contacts, referred to as the Midani 2018 cohort, 20 developed infection during the follow-up period, and 45 remained uninfected (**Figure 1**). Among the 20 contacts who became infected, 10 had no symptoms during the 30-day follow-up period, and were classified as asymptomatic, and 10 developed symptoms (Supplementary Methods). To increase our sample size, we surveyed an expanded cohort (**Table S1a**) by adding 33 samples, including 10 additional pre-infection samples from timepoints of contacts in the Midani 2018 cohort, and 23 samples from 16 newly enrolled contacts from the same place and time (2012-2014, Dhaka, Bangladesh). We used pre-infection samples in order to identify predictive features of disease outcomes in the Midani 2018 cohort, upon which we base the majority of our analyses. We also performed exploratory analyses on the expanded cohort to determine the potential for predictive models to be generalized to larger samples. We used the shotgun metagenomic DNA sequence reads from these samples to characterize four features of the microbiome: 1) relative abundances of microbial species, 2) the presence/absence of sub-species-level strains, 3) metabolic pathway relative abundances, and 4) gene family relative abundances (**Table 1**).

**Table 1.** Assessment of prediction performance for a random forest (RF) model applied to the Midani 2018 and expanded cohorts. Species abundances, strain-specific markers presence/absence, relative abundance of Pfam-grouped gene families, and MetaCyc pathways were used as features. For each dataset, we applied a binary (uninfected vs. infected contacts) and a multi-class (asymptomatic vs. symptomatic vs. uninfected contacts) classifier and reported performance metrics for each dataset. Metrics obtained by the same classifier applied to the same datasets with shuffled class labels (random assignment of labels to samples) are also reported (shuffled). The margins of errors (95% confidence intervals) are reported in parenthesis.

|  |  | RF – Cohort from Midani et al |  |  |  | RF – Expanded |  |  |  |
| --- | --- | --- | --- | --- | --- | --- | --- | --- | --- |
|  |  | Species abundance | Strain markers | Gene families | Pathways | Species abundance | Strain markers | Gene families | Pathways |
| #features |  | 705 | 54953 | 6810 | 443 | 807 | 62965 | 7514 | 461 |
| Infected Vs Uninfected | Accuracy | 0.73<br>(±0.02) | 0.71<br>(±0.02) | 0.76<br>(±0.02) | 0.72<br>(±0.02) | 0.76<br>(±0.03) | 0.69<br>(±0.03) | 0.80<br>(±0.02) | 0.80<br>(±0.03) |
|  | Precision | 0.71<br>(±0.06) | 0.68<br>(±0.06) | 0.77<br>(±0.04) | 0.70<br>(±0.05) | 0.76<br>(±0.03) | 0.70<br>(±0.03) | 0.81<br>(±0.02) | 0.81<br>(±0.03) |
|  | F1 | 0.66<br>(±0.02) | 0.64<br>(±0.03) | 0.71<br>(±0.03) | 0.66<br>(±0.03) | 0.75<br>(±0.03) | 0.68<br>(±0.03) | 0.80<br>(±0.02) | 0.80<br>(±0.03) |
|  | AUC | 0.61<br>(±0.05) | 0.62<br>(±0.04) | 0.74<br>(±0.04) | 0.71<br>(±0.04) | 0.83<br>(±0.02) | 0.76<br>(±0.03) | 0.87<br>(±0.02) | 0.88<br>(±0.02) |
| Shuffled | F1 | 0.55<br>(±0.04) | 0.56<br>(±0.04) | 0.56<br>(±0.04) | 0.56<br>(±0.05) | 0.40<br>(±0.03) | 0.45<br>(±0.03) | 0.48<br>(±0.03) | 0.44<br>(±0.03) |
|  | AUC | 0.40<br>(±0.04) | 0.57<br>(±0.04) | 0.50<br>(±0.05) | 0.50<br>(±0.04) | 0.39<br>(±0.03) | 0.52<br>(±0.03) | 0.51<br>(±0.03) | 0.46<br>(±0.03) |
| Asymptomatic vs Symptomatic vs Uninfected | Accuracy | 0.70<br>(±0.02) | 0.70<br>(±0.02) | 0.69<br>(±0.01) | 0.69<br>(±0.01) | 0.68<br>(±0.01) | 0.60<br>(±0.03) | 0.69<br>(±0.02) | 0.67<br>(±0.03) |
|  | Precision | 0.53<br>(±0.03) | 0.53<br>(±0.03) | 0.60<br>(±0.02) | 0.59<br>(±0.02) | 0.60<br>(±0.02) | 0.53<br>(±0.03) | 0.61<br>(±0.02) | 0.59<br>(±0.02) |
|  | F1 | 0.60<br>(±0.02) | 0.59<br>(±0.02) | 0.57<br>(±0.02) | 0.57<br>(±0.02) | 0.62<br>(±0.02) | 0.55<br>(±0.03) | 0.64<br>(±0.02) | 0.62<br>(±0.02) |
|  | AUC | NA | NA | NA | NA | NA | NA | NA | NA |
| Shuffled | F1 | 0.48<br>(±0.04) | 0.49<br>(±0.04) | 0.46<br>(±0.03) | 0.55<br>(±0.03) | 0.41<br>(±0.03) | 0.35<br>(±0.03) | 0.44<br>(±0.04) | 0.37<br>(±0.03) |
|  | AUC | NA | NA | NA | NA | NA | NA | NA | NA |

### Predicting susceptibility to *V. cholerae* infection with Random Forests

We first used an RF model to predict *V. cholerae* susceptibility (developing infection or remaining uninfected) from baseline microbiome features (**Figure 1**). In the Midani 2018 cohort, functional pathways and gene families predicted infection significantly better than random (Two-sample *t*-tests comparing area under the curve [AUC] across 20 replicate 3 fold cross-validations; *p* < 0.05) compared to data with shuffled (randomized) labels, and also predicted infection better than species or strain features (**Tables 1 and S2**). Pathways and gene families had significantly higher mean AUCs (0.71 and 0.74, respectively) compared to species or strains (0.61 and 0.62) (*p* < 0.05; **Table 1**; **Figure S1, Table S3**).

To determine the minimum number of metagenomic features required for prediction, we repeated the analysis using smaller subsets of features. Using only 30 species, 60 gene families or pathways, or 200 strains achieved similar cross-validation AUC values (**Figure S2**). We then trained an RF model on this reduced number of selected features, yielding improved predictions for all feature types (**Figure S1; Table S4**). This suggests that only a limited number of strains, species, genes and pathways in the gut microbiome at the time of exposure are sufficient to predict *V. cholerae* susceptibility. For example, prediction using strain-level markers after feature selection yielded an AUC of 0.95 (**Table S4**). However, such high AUC values should be treated with caution because the models can be overfit when a supervised feature selection step is applied on the same data used to train the model (18). Because we did not have a fully independent validation cohort (*e*.*g*. from another continent) to test our model, we decided to use the features selected from the Midani cohort to make predictions on the Expanded dataset. Using the same features selected from the Midani 2018 training dataset, we made predictions on the Expanded cohort and achieved AUCs between 0.89 and 0.93 for prediction of infection using the four types of features (**Table S4**). Again, because the expanded cohort partly overlaps with the Midani cohort, and includes some repeated samples from the same individuals over time, these results could also be prone to overfitting, but they demonstrate the potential for generalized predictions.

Finally, we repeated the RF analysis using all features in the expanded dataset, whichh increased predictive performance relative to the original Midani cohort (**Figure S1**). Once again, genes and pathways outperformed species and strains according to all metrics, with AUC reaching ∼0.88 using cellular pathways (**Table 1**). This improvement in the expanded cohort also highlights the importance of using larger, more balanced datasets as input to predictive models.

### Improved prediction compared to known factors impacting susceptibility

To put the metagenomic predictions in context, we compared their predictive power and accuracy to clinical and demographic factors (**Table S1a**). Three of these factors (age, baseline vibriocidal antibodies and blood group) are known to impact susceptibility to *V. cholerae* infection (6,15) and we used them to train RF models (**Table S5**). As expected, contacts who became infected tended to be younger and have lower baseline antibody titers than those who remained uninfected (**Table S1b**), but these small differences were not sufficient to train a significantly predictive model. An RF model trained on the seven clinical and demographic factors did not perform better than a random model with shuffled labels (AUC=0.60, *p*=0.66; **Figure 2**). Predictions were not improved using all species-level metagenomic features present at the time of exposure to *V. cholerae* (AUC=0.61), but significantly improved using a selected number of species (AUC=0.80, *p* < 1 ×10^−7^). The use of all gene families or a selected number of genes showed an increased predictive performance (AUC=0.74 and AUC=0.89 respectively; **Figure 2**) compared to species-level or clinical and demographic contact data (*p* < 1 ×10^-7^ for all comparisons). We again note the caveat that models with selected features may be overfit and represent an upper bound for predictive power. Even without feature selection, we found that gene families clearly provide superior predictions, and adding clinical data did not improve the predictions based on microbiome features alone (**Figure 2**). Together, these results demonstrate that gene families present in the gut microbiome at the time of exposure contain more information about *V. cholerae* susceptibility compared to species-level or clinical and demographic contact data.

**Figure 2.**
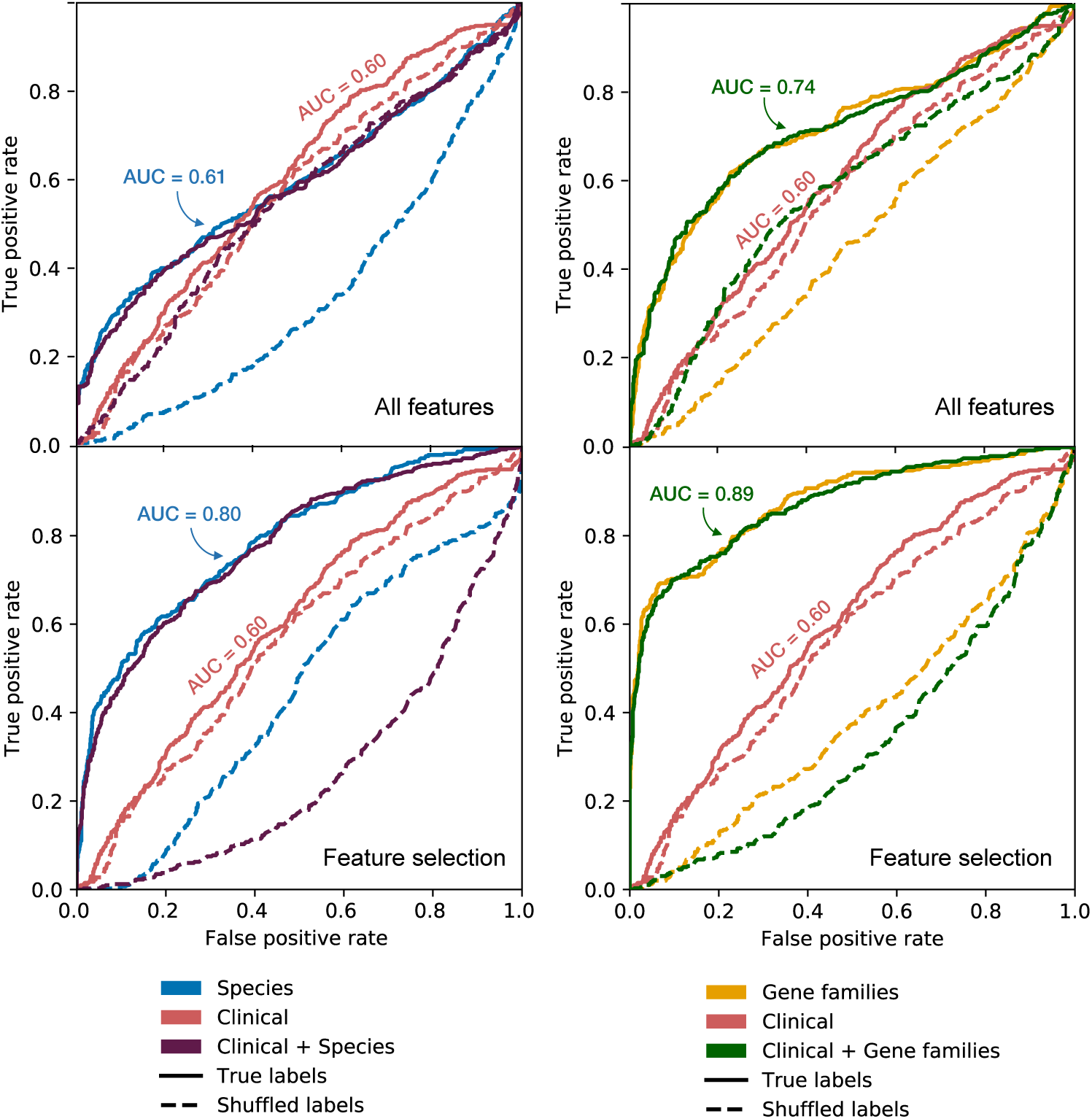
Metagenomic features predict *V. cholerae* infection better than clinical and demographic features. Random forest prediction of infection status was applied to 7 clinical and demographic features, and compared with all species and all gene families (top row), as well as 30 selected species features from metagenomes and 60 selected gene family features (bottom row), or a combination of clinical, demographic and metagenomic features. Plots show receiver operating characteristic (ROC) curves (average across cross-validations) for the Midani 2018 dataset. Shuffled labels represent the prediction run on a dataset with a random assignment of infection outcomes. AUC = area under the curve.

### Disease severity is more difficult to predict than likelihood of infection

To predict symptomatic disease among infected individuals (**Figure 1**), we divided samples into uninfected, symptomatic and asymptomatic groups and again applied the RF approach. We used the F1 score as a performance metric since it is well suited for uneven class distributions in our uninfected/symptomatic/asymptomatic comparison. Applied to the Midani 2018 cohort, this model predicted outcomes significantly better than random (shuffled labels) using species, strains or pathway data, but not gene families (**Table 1;** see **Table S3** for *p*-values). However, the F1 scores for the symptomatic/asymptomatic predictions were systematically lower (mean scores ranging from of 0.57 to 0.60) than for the infected/uninfected prediction (means ranging from 0.64 to 0.71). Using the expanded cohort, the scores were improved only slightly (**Table 1**). These results suggest that disease severity is predictable in principle, but with greater uncertainty than the infection outcome.

### Taxonomic biomarkers of disease susceptibility and severity

Predictive features in the gut microbiome identified to a species/strain or gene level allow the possibility of experimental follow-up to investigate mechanisms of the associations we observed. We characterized the most predictive species, pathways, and gene families (**Tables S6-S9**). The most common discriminating species in individuals that remained uninfected during the follow-up period were *Eubacterium rectale, Campylobacter hominis, Ruminococcus gnavus, Bacteroides vulgatus, Veillonella parvula* and members of the *Prevotella* and *Eubacterium* genera (**Figures 3A, S3A and S4A**). These species are ranked by their importance score, which is effectively their relative weighting in the RF model. Several species associated with contacts that developed *V. cholerae* infection belonged to the genera *Bifidobacterium, Actinomyces* or *Collinsella*, and many of the species were also associated with asymptomatic infection (**Figures 3B, S3B and S4B**), including three species of *Bifidobacterium*. The top predictive species in contacts who developed symptomatic infection were *Clostridium ventriculi* (formerly *Sarcina ventriculi*), *Streptococcus parasanguinis* and members of *Veillonella. Shigella* species were also associated with the gut microbiome of persons who developed symptomatic *V. cholerae* infection, although persons enrolled in this study were *Shigella* stool-culture negative. *Shigella* identified by DNA presence in stool may be the result of recent or resolving infection, or may be present at subclinical levels due to ingestion of contaminated water. The features identified by the multivariate RF model were confirmed using univariate statistics for the uninfected/infected prediction (**Figure S5)**, but the overlap was poorer for the uninfected/symptomatic/asymptomatic prediction (**Figure S6)**. This is consistent with the difficulty of predicting disease severity.

**Figure 3.**
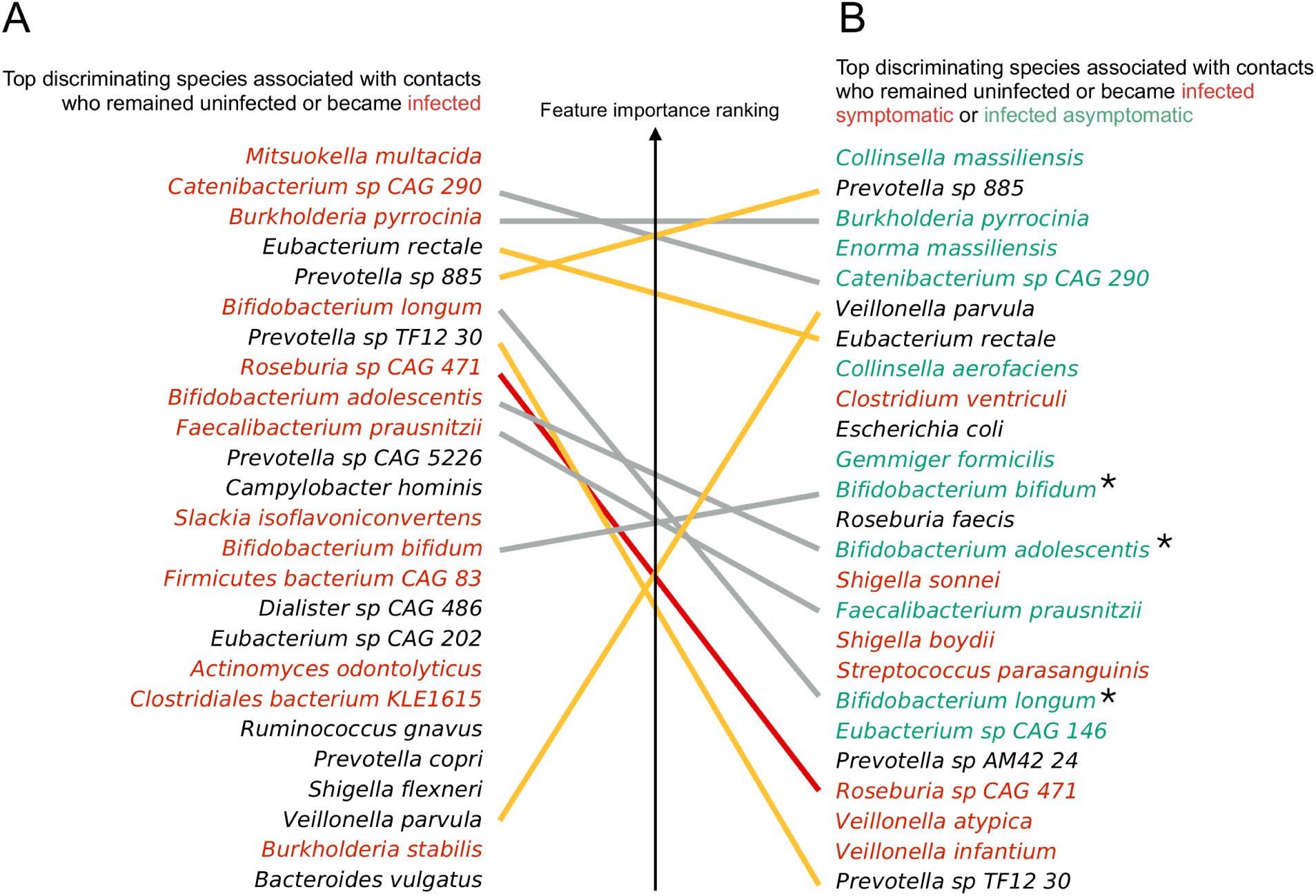
Most important discriminating species of the gut microbiome at the time of exposure to *V. cholerae* identified in the Midani 2018 dataset, classified by clinical outcome. (A) Species associated with contacts that became infected (red) or remained uninfected (black) during follow-up. (B) Species associated with contacts who remained uninfected (black), or became infected asymptomatic (green), or symptomatic (red) during follow-up. The top 25 most important features for discriminating between classes in the RF model are shown here; see Table S6 for the full list. Yellow lines connect species associated with uninfected individuals in both (A) and (B); red lines connect species associated with infection in (A) and symptomatic disease in (B); grey lines connect species associated with infection in (A) but asymptomatic infection in (B). Three species of *Bifidobacterium* are marked with an asterisk.

In general, the most important species were selected by the model because of differences in relative abundance at baseline among uninfected/symptomatic/asymptomatic outcomes (**Figure S7, S8**). In rare cases, species presence/absence information was predictive. For example, *Ruminococcus gnavus*, is absent (near or below limit of detection) in most of the individuals who become infected, but present in many (but not all) of those who remain uninfected (**Figure S7**). Thus, there is no single, strong predictor of infection outcomes, but rather a probabilistic combination of many species, each of relatively modest predictive value.

### Identification of functional biomarkers of disease susceptibility and severity

We also identified gene families in the gut microbiome of persons who remained uninfected during follow-up (**Figures S9 and S10)**, with some of the top gene families involved in DNA repair, transmembrane transporter activity, iron metabolism (indicated with asterisks in **Figure 4**), and genes of unknown function (**Table S8**). Long-chain fatty acid biosynthesis pathways (*e*.*g*. cis-vaccenate, gondoate and stearate) were associated with individuals who remained uninfected, while amino acid biosynthesis and catabolic pathways were associated with individuals who developed infection (**Figures S11 and S12, Table S9**). We identified three iron-related genes associated with remaining uninfected: (1) the ferric uptake regulator Fur, a major regulator of iron homeostasis, (2) thioredoxin, a redox protein involved in adaptation to oxidative and iron-deficiency stress, and (3) the TonB/ExbD/TolQR system, a ferric chelate transporter (19-21). In individuals who became infected but asymptomatic, two genes involved in the conversion of riboflavin into catalytically active cofactors, the riboflavin kinase and the FAD synthetase, were found as the first and the third most discriminant features (**Figure 4, Table S8**).

**Figure 4.**
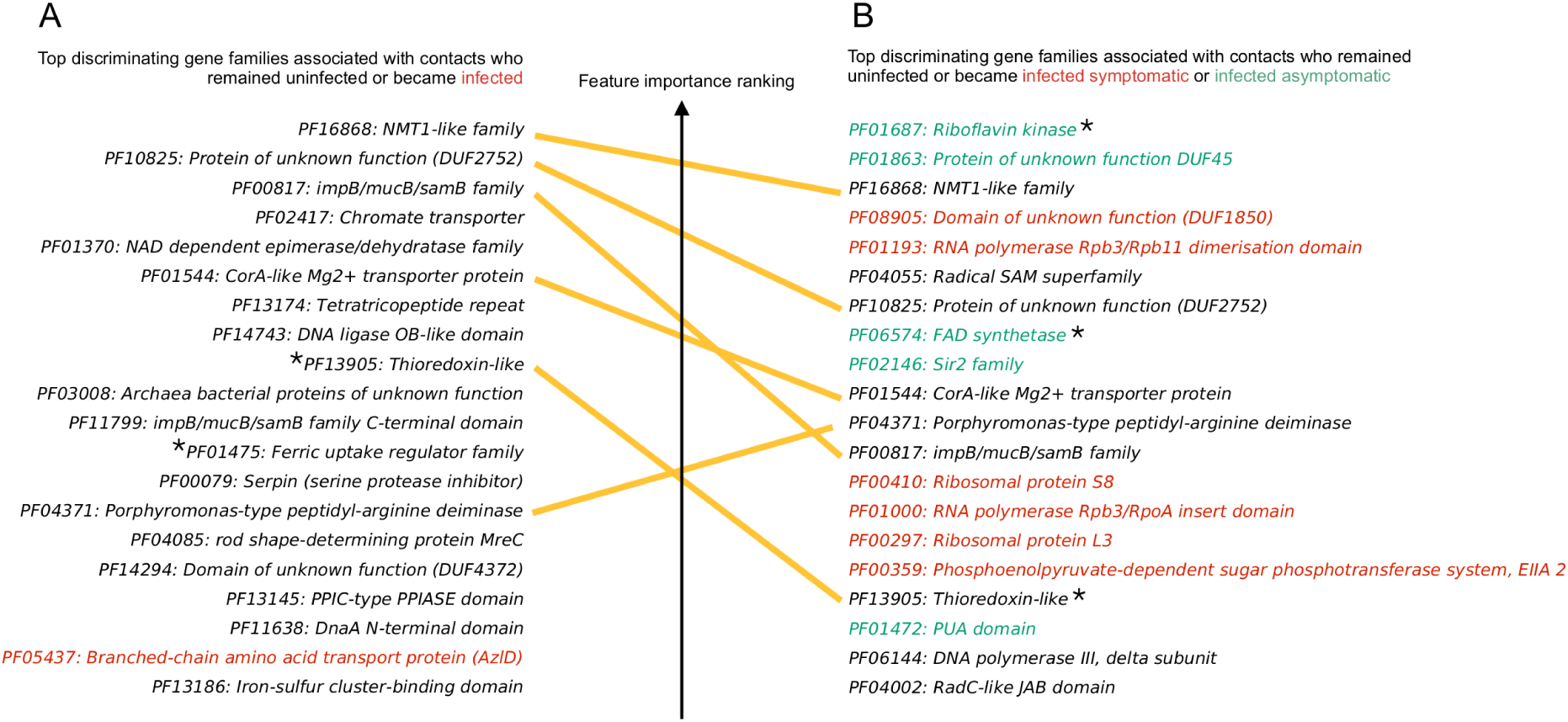
Most important discriminating gene families of the gut microbiome at the time of exposure to *V. cholerae* identified in the Midani 2018 dataset, classified by clinical outcome. (A) Genes families associated with contacts that became infected (red) or remained uninfected (black) during follow-up. (B) Genes families associated with contacts who remained uninfected (black), or became infected asymptomatic (green), or symptomatic (red) during follow-up. The top 25 most important features for discriminating between classes in the RF model are shown here; see Table S8 for the full list. Yellow lines connect species associated with uninfected individuals in both (A) and (B). Asterisks indicate genes involved in redox or iron metabolism.

We next asked which taxa in the microbiome likely encoded these genes. In some cases, specific taxonomic groups corresponded to discrete gene functions. For example, several iron metabolism-related gene families tend to be encoded by *Prevotella* genomes (**Figure S14**). In other cases, the major contributors to protective gene families were unclassified (**Figures 5 and S13**). These results partly explain why gene families or pathway features tend to outperform species-level features in predicting infection status – because predictive gene families are distributed across many species, including several with poor taxonomic annotation or families lacking representation in taxonomic databases.

**Figure 5.**
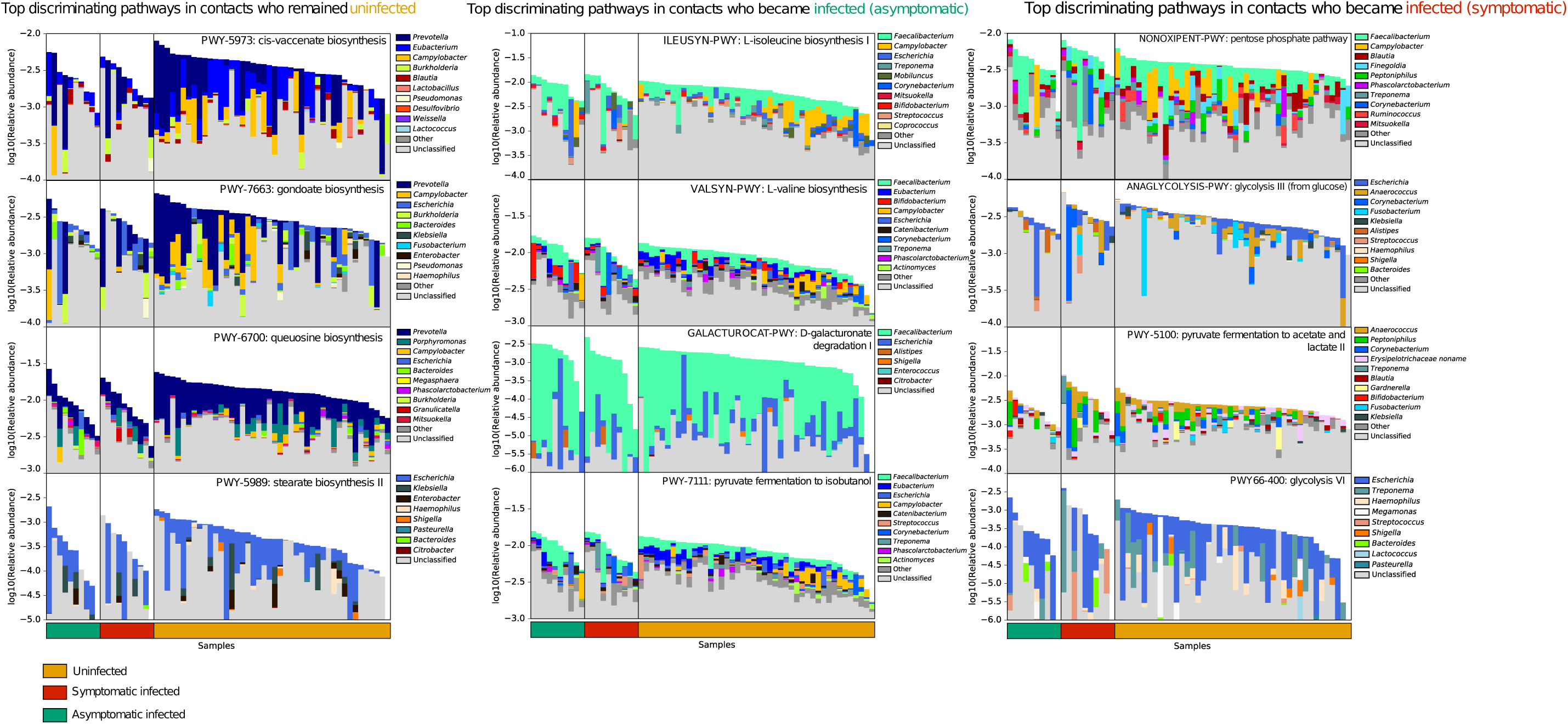
Top predictive cellular pathways of the gut microbiome at the time of exposure to *V. cholerae* in the Midani 2018 cohort, annotated by their taxonomic contributors. The four top-ranked pathways associated with uninfected contacts (left column), contacts who developed asymptomatic infection (middle), and contacts who developed symptomatic infection (right column) are shown. Total bar height reflects log10-scaled community relative abundance of each pathways. The contributions of each genus to encoding these pathways are shown as stacked colors within each bar, linearly scaled within the total. See Table S9 for the complete list of pathways

## DISCUSSION

The gut microbiome is a potentially modifiable host risk factor for cholera, and identification of specific genes and strains correlated with susceptibility is needed for experimental testing to understand the mechanisms of observed correlations. Compared to a previous study using a single marker gene, shotgun metagenomics provides this degree of resolution, potentially to the species and strain level, and to the level of individual genes and cellular functions. We found that gene families in the gut microbiome at the time of exposure to *V. cholerae* were more predictive of susceptibility compared to taxonomic or clinical and demographic information. Selecting a subset of the most informative features further improved predictions, but using these selected features may lead to overfitting. This suggests an upper limit to predictive power that requires validation in larger, independent cohorts.

Most of the top predictive biomarkers were associated with remaining uninfected after exposure to *V. cholerae*. An example is the genus *Prevotella*, including several strains within *Prevotella* sp. 885, identified only at the genus level in a previous study (15). *Prevotella* species are hypothesized to be beneficial members of the microbiota in healthy individuals in non-Westernized countries, and this species is a potential candidate for follow-up experimental studies in *V. cholerae* susceptibility (14,22,23).

Several species known to ferment mucin glycans into short chain fatty acids (SCFAs) correlated with remaining uninfected, including *Eubacterium rectale, Ruminococcus gnavus* and *Bacteroides vulgatu*s (24,25). This finding is consistent with experiments of SCFAs applied to animal models. *B. vulgatus* has been shown to inhibit *V. cholerae* colonization in mice, an effect that was dependent upon SCFA production (13). SCFAs are known to impact immune cell development and attenuate inflammation by inhibiting histone deacetylases and other mechanisms of altering gene expression (26-29).

All three *Bifidobacterium* species associated with contacts that developed infection were also associated with asymptomatic rather than symptomatic disease, and prior work on this genus supports several hypotheses for this relationship. First, *Bifidobacteria* are known to produce the SCFA acetate that can protect against enteric infection in mice (30,33,34). SCFAs are also known to inhibit cholera toxin-related chloride secretion in the mouse gut, reducing water and sodium loss, and have been observed to increase cholera toxin-specific antibody responses (31-33). *Bifidobacteria* are also major producers of lactate, a metabolite that has been shown to impair *V. cholerae* biofilm formation, a function that can impact virulence (12). Lastly, *B. bifidum* and *B. adolescentis* are known to reduce the activity of *V. cholerae* type VI secretion systems through modification of bile acids (9).

Metagenomics also allowed us to identify bacterial functions that could impact the ability of *V. cholerae* to compete and colonize the gut. For example, several gene families involved in iron transport, iron regulation, and riboflavin conversion appeared among the top twenty features associated with uninfected and asymptomatic individuals, suggesting that competition for iron might be a protective mechanism of the gut microbiota against *V. cholerae*, as in other pathogens (7). Iron is often a limiting redox cofactor in the gut, and bacteria have evolved strategies to solubilize and internalize iron (34,35). Riboflavin (another major redox cofactor in bacteria) and iron levels are reciprocally regulated in *V. cholerae*, and riboflavin may allow *V. cholerae* to overcome iron limitation in the gut (34,36). A gut microbiota more competitive for iron could therefore help resist *V. cholerae* colonization or reduce its virulence. Further work is thus needed to understand mechanisms of how the enrichment of these genes may protect people after exposure to *V. cholerae*.

Our results are currently not generalizable beyond the study cohort in Dhaka, Bangladesh, since a similar cohort in another geographic location is not available. As with any association-based study (37), it is unknown if any of the metagenomic features that correlate with protection from *V. cholerae* infection are causal, and many may be markers of clinical or environmental factors that themselves impact susceptibility. Despite our deep sequencing and collection of standard cholera risk factors, our study was unable to measure all potentially relevant environmental or clinical risk factors. In line with recent studies in Dhaka, we assume that *V. cholerae* transmission occurs mainly within households (3) and did not consider how the mode of transmission (*e*.*g*. waterborne or not) might affect outcomes. It has also been noted that microbiome-disease associations may be poorly portable across human populations (37). For instance, we identified species of *Prevotella* as protective features in Bangladesh, but *Prevotella* is much less abundant and less diverse in Western countries (22). It thus remains to be seen if protective gene features (*e*.*g*. iron metabolism) are encoded in other species of the microbiome outside endemic areas like Bangladesh, or if people outside these areas are simply at greater risk for cholera. Further experimental characterization of metagenomic features correlated with protection from infection or symptoms are needed to understand if factors we identified impact *V. cholerae* pathogenesis or host responses to infection. Ultimately, the strains and functionalities identified have the potential to inform microbiota-based therapeutics to ameliorate or prevent disease. Our results show the power of metagenomic data from the gut microbiome to predict health outcomes such as susceptibility to infection and disease severity.

## ACKNOWLEDGEMENTS

We thank Meti Debela for technical assistance. Finally, we are grateful to the people of Dhaka where our study was undertaken; to the field, laboratory and data management staff who provided tremendous effort to make the study successful; and to the people who provided valuable support in our study. The icddr,b gratefully acknowledges the Government of the People’s Republic of Bangladesh; Global Affairs Canada (GAC); Swedish International Development Cooperation Agency (Sida) and the Department for International Development, (UKAid). We declare that we have no competing financial interest.

## ETHICAL STATEMENT

The Ethical and Research Review Committees of the icddr,b and the Institutional Review Board of MGH reviewed the study. All adult subjects provided informed consent and parents/guardians of children provided informed consent. Informed consent was written.

## CONFLICTS OF INTEREST

The authors declare that there are no conflicts of interest.

### Supplementary tables S1-S9 are available at

https://figshare.com/articles/Supplementary_Tables_-_Levade_et_al_2020/12440417

