## Supplementary Methods and Figures for "Predicting *Vibrio cholerae* infection and disease severity using metagenomics in a prospective cohort study"

### Clinical outcomes and metadata collection

All study participants lived in Dhaka city, Bangladesh. Multiple household contacts could be enrolled from one household, and only one index case was enrolled per household. Household contacts were defined as individuals who shared the same cooking pot with an index patient for three or more days. Rectal swab samples were obtained from contacts beginning the day after enrollment of the index case. During the follow up period for contacts, two sets of rectal swabs were collected. The first set were collected during daily home visits, when report of symptoms was also collected, and was used for *V. cholerae* culture.  This data was used for the classification of clinical outcomes. The second set of rectal swabs collected over the first ten days, and day 30, also at the home visit, were used for DNA extraction to conduct the microbiome analyses. Infection status was determined using the first set of rectal swab samples, serologic data from the blood draw, and report of symptoms collected during the observation period. Blood samples collected during the same days from contacts were drawn at the International Center for Diarrheal Disease Research, Bangladesh, in Dhaka, Bangladesh. Symptomatic *V. cholerae* infection was defined by a contact reporting diarrhea within 3 days of a rectal swab positive for *V. cholerae* during the follow-up home visits, or by a four-fold increase in vibriocidal titer with diarrhea during the follow-up period. Two symptomatic contacts were rectal swab negative for *V. cholerae* but showed a four-fold increase in vibriocidal titer and reported diarrhea, and then were determined to be infected based on our inclusion criteria. *V. cholerae* infection (rectal swab positive) was defined as asymptomatic if no diarrhea was reported. All contacts in our study that developed infection had had the same serotype of *V. cholerae* as the infected household case (either Inaba or Ogawa).

### Midani and Expanded cohort description

We used metagenomic reads from 65 of the 76 contacts from a previously published study by our collaborators, and these contacts are referred to as the Midani 2018 cohort (1). Eleven of the contacts from this study had insufficient DNA to perform metagenomic sequencing. Each sample from this cohort was sampled on day 2, the day of the enrollment of household contacts, one day after the presentation of the household index case at the hospital. To minimize the chance that the contact acquired a *V. cholerae* infection between index case enrollment and the first contact sample, the first household contact sample was always taken within a day, and generally <24h since the enrollment of the index case. To increase power in our study, we added household contacts samples from additional households, and also added samples from additional time points of Midani 2018 cohort contacts (see Table S1). This larger group of samples is referred to as the Expanded cohort, and is comprised of the 65 samples from the Midani 2018 cohort plus 33 additional samples from 16 infected contacts (Figure 1). The 16 infected new contacts in the expanded cohort have multiple samples from different days that are prior to when infection occurred, and therefore have multiple "pre-infection" samples (Table S1). Enrollment and sample collection were identical for the two groups of samples collected. In total, 129 household contacts were initially enrolled. Eighteen contacts were excluded due to a positive rectal swab at the enrollment evaluation, and 3 were excluded due to recent antibiotic use. Nine contacts reported ambiguous clinical outcomes based on clinical and history evaluation (i.e. report of diarrhea with no serologic or culture evidence of *V. cholerae* infection). Six additional enrollees were excluded due to report of diarrhea during the week prior to enrollment. Among the remaining contacts, 10 resulted with DNA evidence of *V. cholerae* infection despite a rectal swab culture negative for *V. cholerae* at the time of enrollment. Lastly, 13 contacts were excluded due to failure to amplify DNA from rectal swabs or sequencing failure.

### DNA extraction and sequencing

Fecal samples and rectal swabs were collected during home visit and immediately placed on ice and stored at -80°C until DNA extraction. For each sample, DNA was extracted as previously described [4](1). Briefly, samples were processed using PowerSoil DNA extraction kits (Qiagen) after pre-heating to 65°C for 10 min and to 95°C for 10 min. A total of 98 samples were selected for library construction with NEBNext Ultra II DNA library prep kit and sequenced on the Illumina HiSeq 2500 (paired-end 125 bp) and the Illumina NovaSeq 6000 S4 (paired-end 150 bp) at the Genome Québec sequencing platform (McGill University).

### Sequence preprocessing

Sequencing fastq files were quality checked with FastQC (https://www.bioinformatics.babraham.ac.uk/projects/fastqc/). For removal of human and technical contaminant DNA, metagenomic shotgun sequences were aligned to the PhiX genome and the human genome (hg19) with Bowtie2 (2). Host-decontaminated fastq files were then quality filtered using Trimmomatic, a computational tool for removing low quality data (3), discarding all reads shorter than 75 nucleotides and with quality Phred score less than 20.

### Taxonomic and functional profiling

We performed taxonomic profiling of archaea, bacteria and microbial eukaryotes using MetaPhlAn2 (version 2.9) (4), with default parameters. Briefly, MetaPhlAn2 maps shotgun metagenomic sequencing reads against a precomputed database of clade-specific markers to produce a robust estimate of the taxonomic clades present in the microbiome sample and estimate their relative abundance. In addition to species-level relative abundance, we used MetaPhlAn2 to characterize the presence/absence of strain-specific markers for each sample. Species abundances are real numbers in the range [0,1] while markers assume binary values.

Species-resolved functional profiling was completed using HMP Unified Metabolic Analysis Network 2 (HUMAnN2, version 2.8.1) (5), which maps reads to a sample-specific reference database from the pangenomes of the subset of species detected in the sample by MetaPhlAn2, quantifying gene presence and abundance on a per-species basis. A translated search is then performed against a UniRef-based protein sequence catalog for reads that fail to map to one of the detected species. The results are abundance profiles of gene families (UniRef90s) stratified by species contributing to those genes, which can be regrouped in higher level grouping categories (Pfam domains in this data set). Finally, gene families were further analyzed to reconstruct metabolic pathways abundance according to the MetaCyc pathway catalog (5).

**Statistical analyses**

Univariate analyses was performed to identify discriminatory biomarkers (species, protein families and pathways) using linear discriminant analysis (LDA) effect sizes (LEfSe) (6). LEfSe first uses the non-parametric factorial Kruskal-Wallis test to detect features with significant differential abundance for each class of interest, then uses LDA to estimate the effect size of each differential abundant feature (6). In our case, the significance threshold was set at $p$=0.05 and an LDA effect size of 2 was used for discriminative features.

**Random-forest based machine learning approach**

MetAML (7) was applied to four types of quantitative profiles: taxonomic species-level relative abundances, strain-specific markers presence/absence patterns, MetaCyc pathways and gene-families grouped by Pfam domains relative abundances. In each case, we used Random Forest (RF) as classifier and set the following parameters as previously described by (8): the number of estimator trees was equal to 1000 and the quality of split at each tree node was evaluated using Shannon entropy. The minimum number of samples per leaf and the number of features per tree were respectively set as 1 and 30%, except for the strain-specific markers presence/absence profile, where the number of features equal the square root of the total number of features, due to the higher number of features. One of the advantages of RF models is that they can intrinsically integrate binary and multi-class classification problems, and implicitly provide a list of the most informative features to the predictive model. Another advantage of RFs is the built-in feature selection step during the model generation phase, which allows a selection of a reduced subset of the most important features for discriminating between classes. Adding a feature selection step to a model is a useful way to remove irrelevant features, especially in datasets with high dimensionality. Feature selection can also reduce overall training time and the risk of overfitting (7).

The feature relative importance values were used to perform an embedded feature selection strategy, implemented as described in (7), in addition to the RF model, for two (Uninfected vs Infected contacts) and three classes (Asymptomatic infected vs Symptomatic infected vs Uninfected contacts). Feature selection benefits include removal of irrelevant features, especially in datasets with high dimensionality, to decrease the overall training time and reduce the risk of model overfitting (9). Each cohort (the Midani_2018 and the expanded cohort) was analyzed independently for the RF model, using a stratified cross-validation approach. In the feature selection model, we chose the minimum number of features that maximized the accuracy in order to generate a final model on this limited number of features (Figure S2). The same set of selected features determined by the Midani_2018 dataset was used for the two cohorts in this approach. In all cases, the sensitivity and accuracy of the models generated were tested by performing stratified 3-fold cross validations, repeated and averaged on 20 independent runs. Finally, the results obtained for the original classification problem were compared with those obtained by a random classifier (denoted in the paper as Shuffled). For this purpose, we shuffled randomly the labels of all the samples, and used these same settings. Difference of performance between classifiers and statistical significance were calculated as described in Pasolli *et al* (7).

The following performance metrics were reported: 1) the overall accuracy (i.e., the proportion of outcomes correctly predicted), 2) the precision (i.e., the number of correct positive samples divided by the number of samples predicted as positive), 3) the recall (i.e. the true positive rate, or power), and 4) the F1 score, which is the harmonic mean of precision and recall. For binary classification problems, class posterior probabilities were used to plot the Receiver Operating Characteristic (ROC) curve, which represents the true positive rate (i.e., the recall) against the false positive rate (i.e., the number of wrong positive samples divided by the total number of non-positive samples). From the ROC curve, the program computes the area under the curve (AUC), which can be interpreted as the probability that a randomly selected positive sample will have a higher classification result than a randomly selected negative one. The AUC ranges in [0, 1], where 0.5 corresponds to random prediction.

### Supplementary Figures

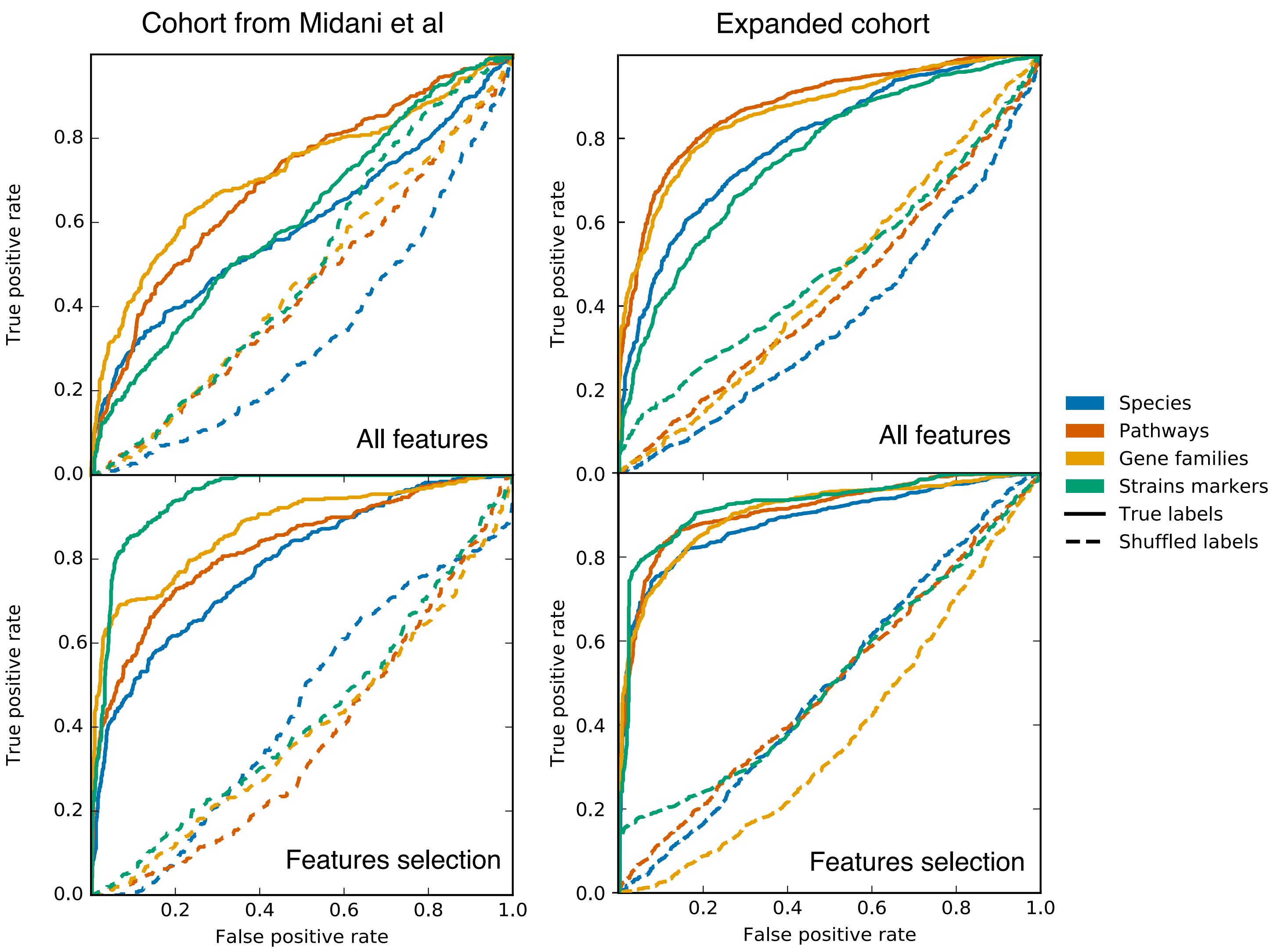


**Figure S1. Classification result from species abundance, strain-level markers presence/absence, pathways and gene family abundances.** Four different classifications of microbiome information were used to predict disease susceptibility (Uninfected vs Infected) using a random forest algorithm and stratified 3-fold cross validation. Plots show Average ROC curves by using the four type of features for two different datasets (the Midani et al data and the expanded dataset). Random Forest was applied on all features (top row) and on a set of selected features (bottom row).

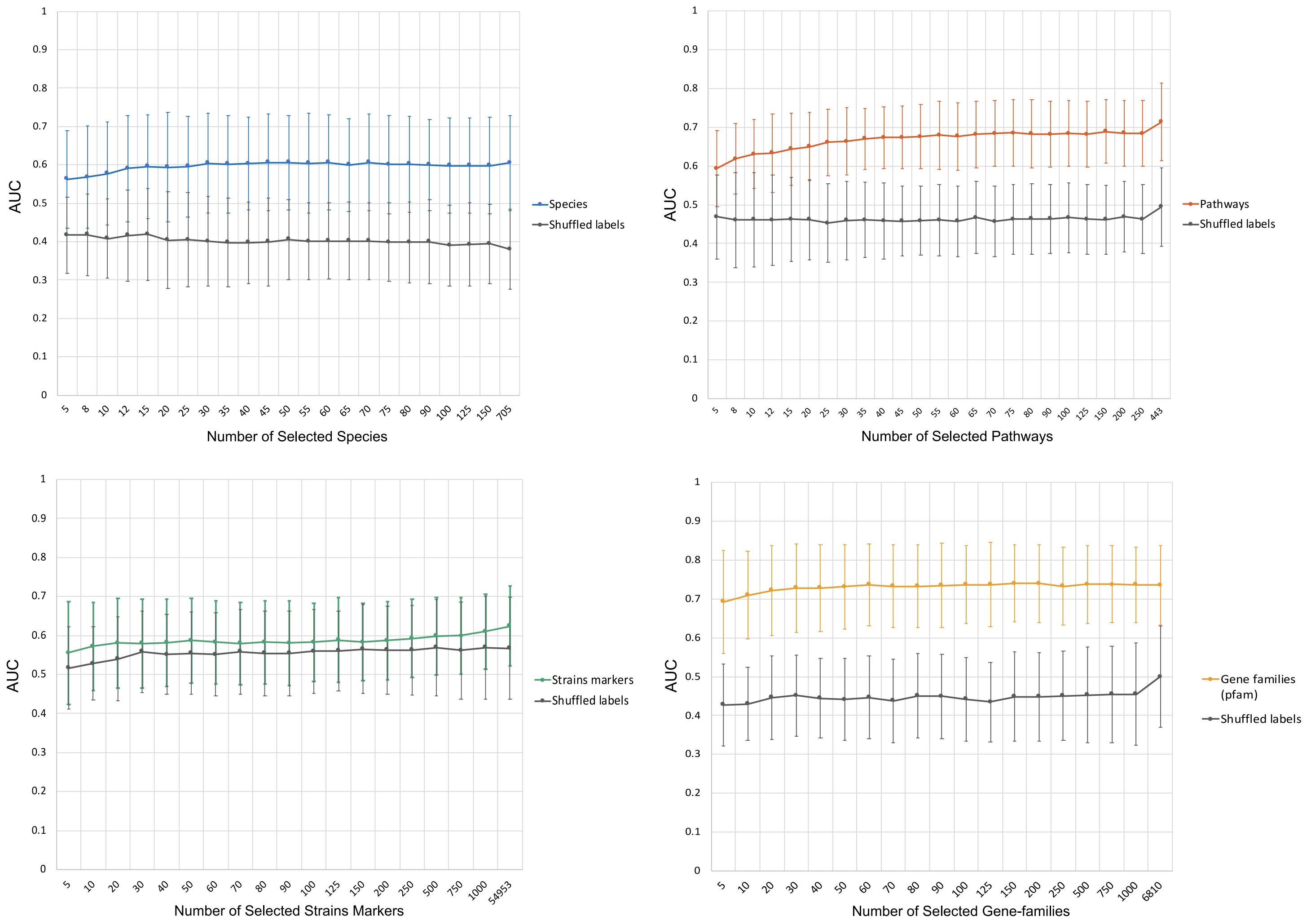


**Fig. S2. Identification of a minimal number of features for each type of biomarker.** Prediction performances (AUC values) at increasing number of microbial species, pathways, strains markers and genes families, obtained by retraining the random forest model on the top-ranking features identified with a first random forest model training with a 3-fold cross-validation approach.

A

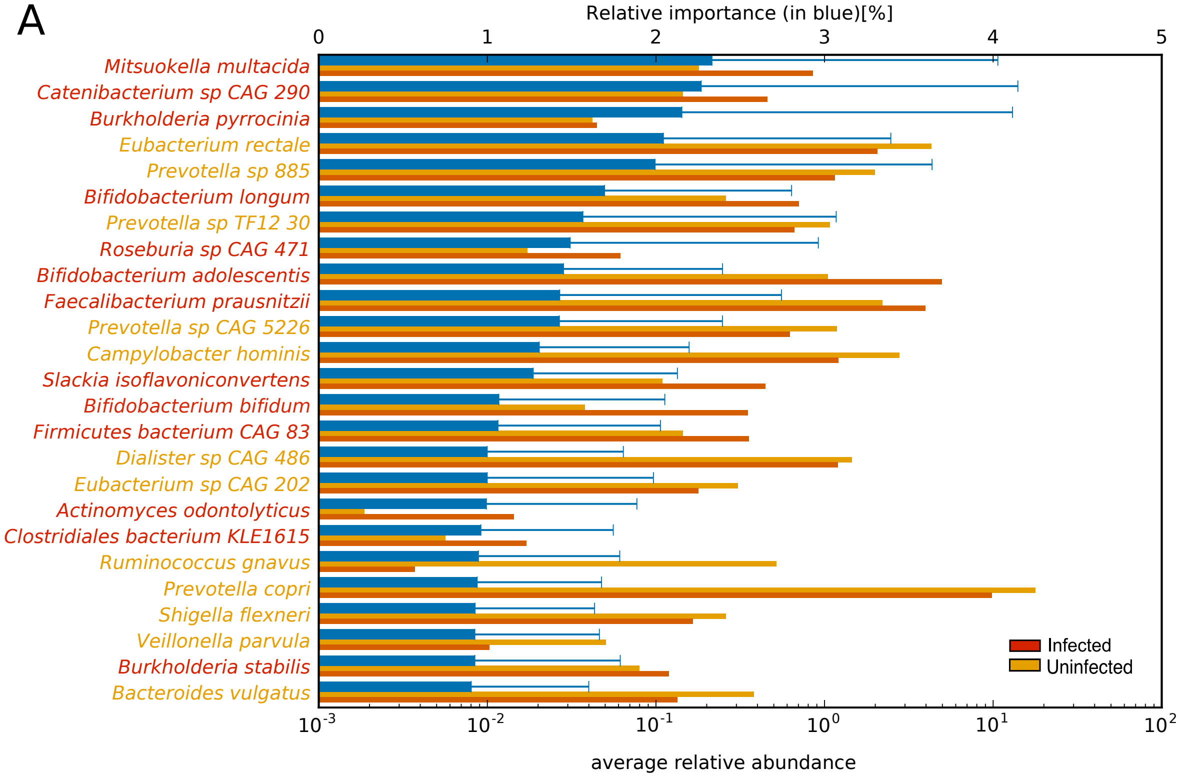
Relative importance (in blue)[%]
0
1
2
3
4
5
Mitsuokella multacida
Catenibacterium sp CAG 290
Burkholderia pyrrocinia
Eubacterium rectale
Prevotella sp 885
Bifidobacterium longum
Prevotella sp TF12 30
Roseburia sp CAG 471
Bifidobacterium adolescentis
Faecalibacterium prausnitzii
Prevotella sp CAG 5226
Campylobacter hominis
Slackia isoflavoniconvertens
Bifidobacterium bifidum
Firmicutes bacterium CAG 83
Dialister sp CAG 486
Eubacterium sp CAG 202
Actinomyces odontolyticus
Clostridiales bacterium KLE1615
Ruminococcus gnavus
Prevotella copri
Shigella flexneri
Veillonella parvula
Burkholderia stabilis
Bacteroides vulgatus
Infected
Uninfected
$10^{-3}$
$10^{-2}$
$10^{-1}$
$10^{0}$
$10^{1}$
$10^{2}$
average relative abundance

B

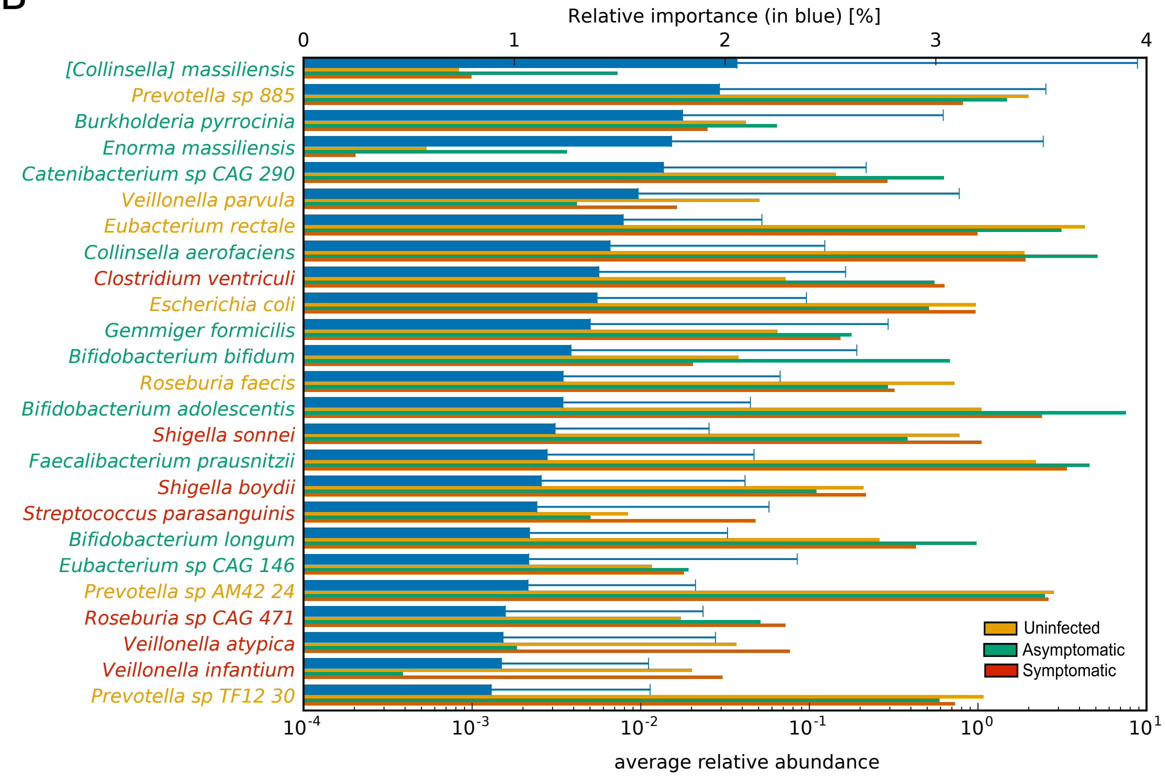
Relative importance (in blue) [%]
0
1
2
3
4
[Collinsella] massiliensis
Prevotella sp 885
Burkholderia pyrrocinia
Enorma massiliensis
Catenibacterium sp CAG 290
Veillonella parvula
Eubacterium rectale
Collinsella aerofaciens
Clostridium ventriculi
Escherichia coli
Gemmiger formicilis
Bifidobacterium bifidum
Roseburia faecis
Bifidobacterium adolescentis
Shigella sonnei
Faecalibacterium prausnitzii
Shigella boydii
Streptococcus parasanguinis
Bifidobacterium longum
Eubacterium sp CAG 146
Prevotella sp AM42 24
Roseburia sp CAG 471
Veillonella atypica
Veillonella infantium
Prevotella sp TF12 30
Uninfected
Asymptomatic
Symptomatic
$10^{-4}$
$10^{-3}$
$10^{-2}$
$10^{-1}$
$10^{0}$
$10^{1}$
average relative abundance

**Fig. S3. Most important discriminating species of the gut microbiome at the time of exposure to *V. cholerae* identified in the Midani 2018 dataset, classified by clinical outcome.** (A) Species associated with contacts that became infected (or remained uninfected) during follow-up. (B) Species associated with contacts that became uninfected/asymptomatic/symptomatic during follow-up. For each species reported on the vertical axis, the top bar (blue) corresponds to the species relative importance (with standard deviation) and the other bars refer to the average relative abundance. The top 25 most important features are shown here; See Table S6 for the full list. Feature relative importance was computed using the Mean Decrease in Impurity strategy, as described in the Methods.

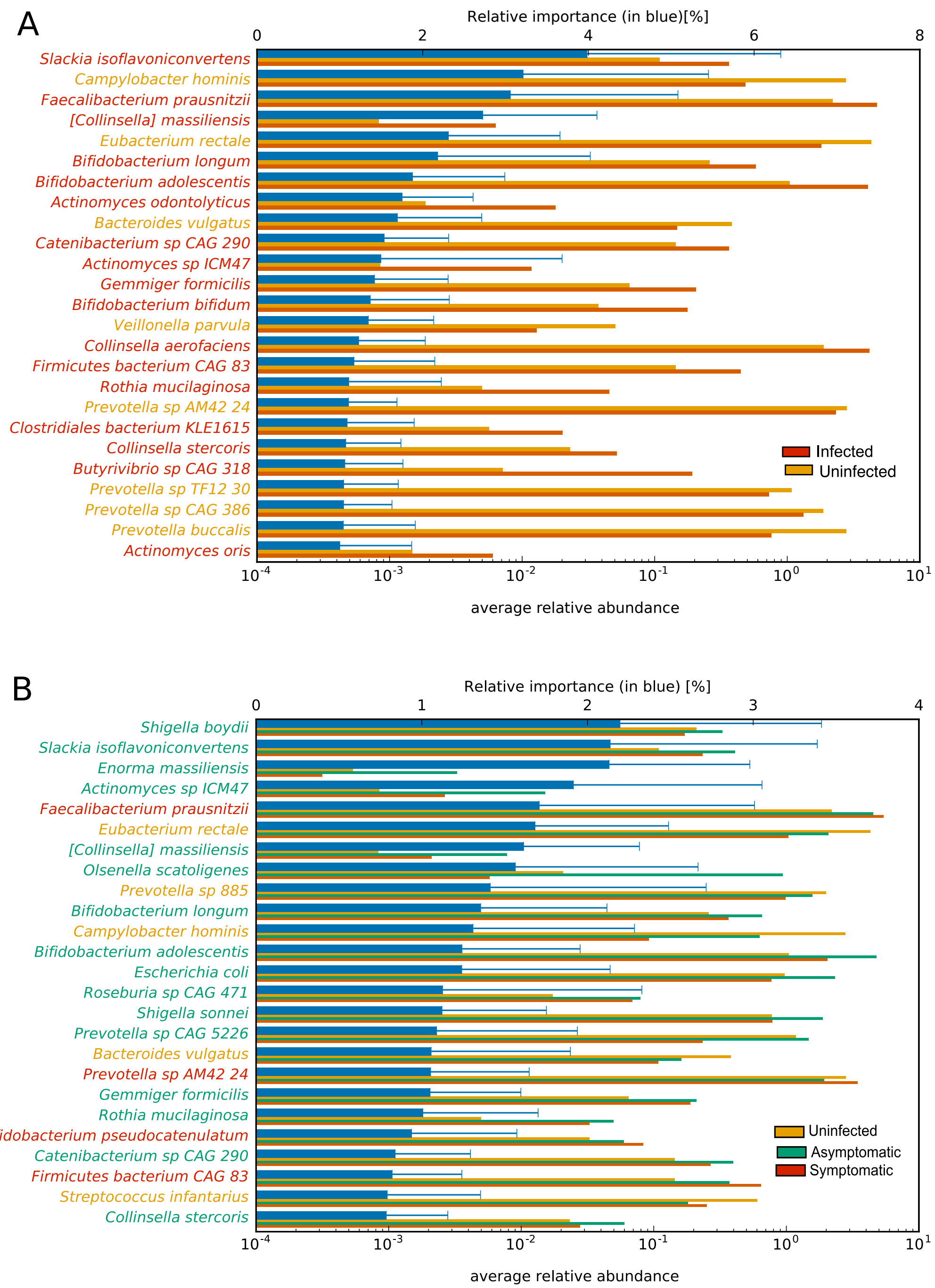


**Fig. S4. Most important discriminating species identified with the random forest algorithm on the expanded dataset for (A) contacts that became infected or remained uninfected and (B) contacts that remained uninfected, or became infected and were asymptomatic vs symptomatic.** For each species reported on the vertical axis, the top bar (in blue) corresponds to the feature relative importance (with standard deviation) and the other bars refer to the average relative abundance. Feature relative importance (blue) is computed using the mean decrease impurity strategy.

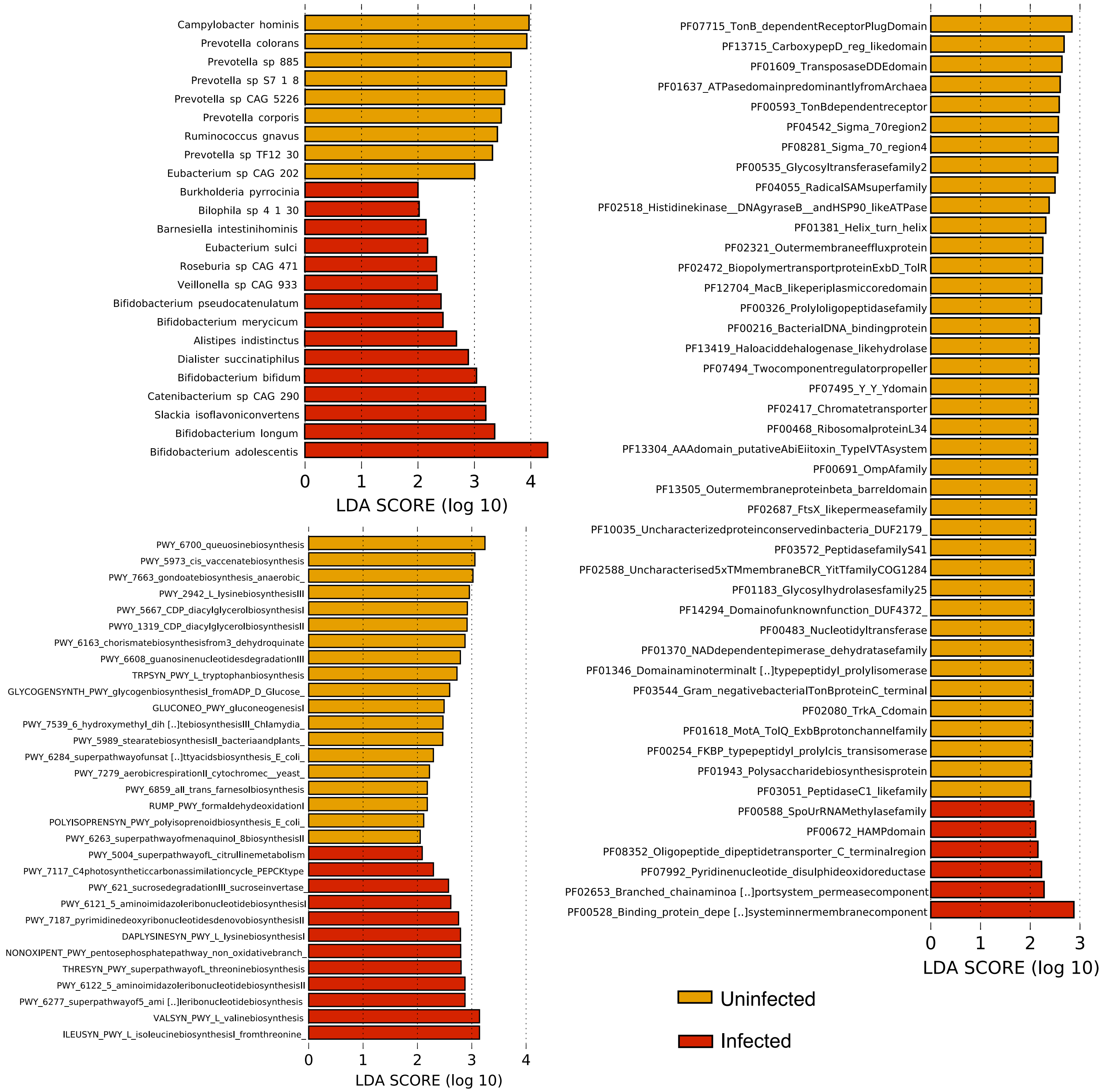

**Fig. S5. Linear discriminant analysis (LDA) scores computed for differentially abundant species, pathways and genes families in the fecal microbiomes of samples from the Midani_2018 cohort for 2 categories.** The cohort is divided in two categories: controls who remained uninfected (yellow) and controls who became infected (red). Length of the bar indicates effect size associated with a feature. $p = 0.05$ for the Kruskal-Wallis H statistic; features with LDA score > 2 are shown.

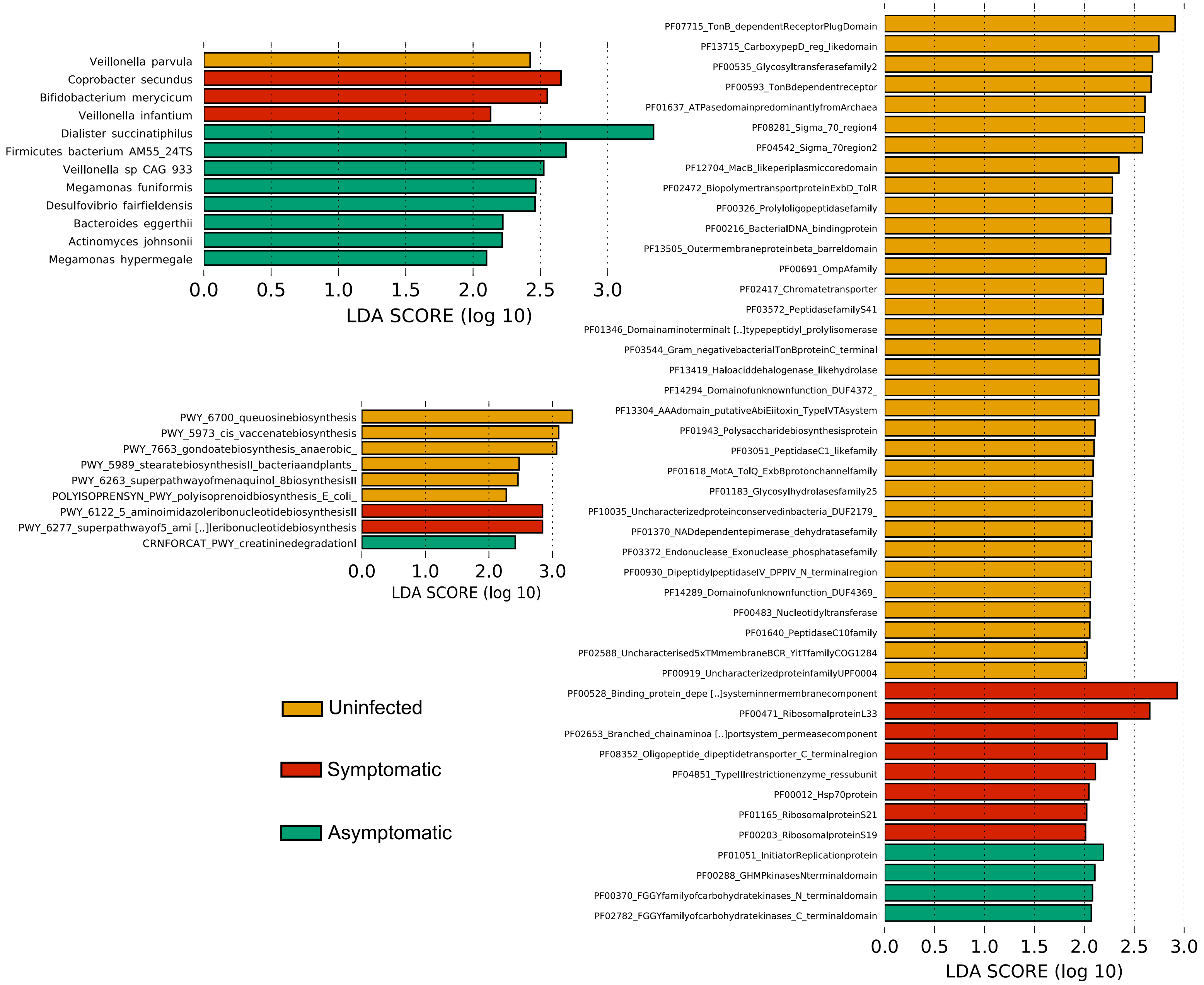


**Fig. S6. Linear discriminant analysis (LDA) scores computed for differentially abundant species, pathways and genes families in the fecal microbiomes of samples from the Midani_2018 cohort for 3 categories.** The cohort is divides in three categories: controls who remained uninfected (yellow), controls who became infected and symptomatic (red), and controls who became infected but stayed asymptomatic (green). Length indicates effect size associated with a feature. $p = 0.05$ for the Kruskal-Wallis H statistic; features with LDA score > 2 are shown.

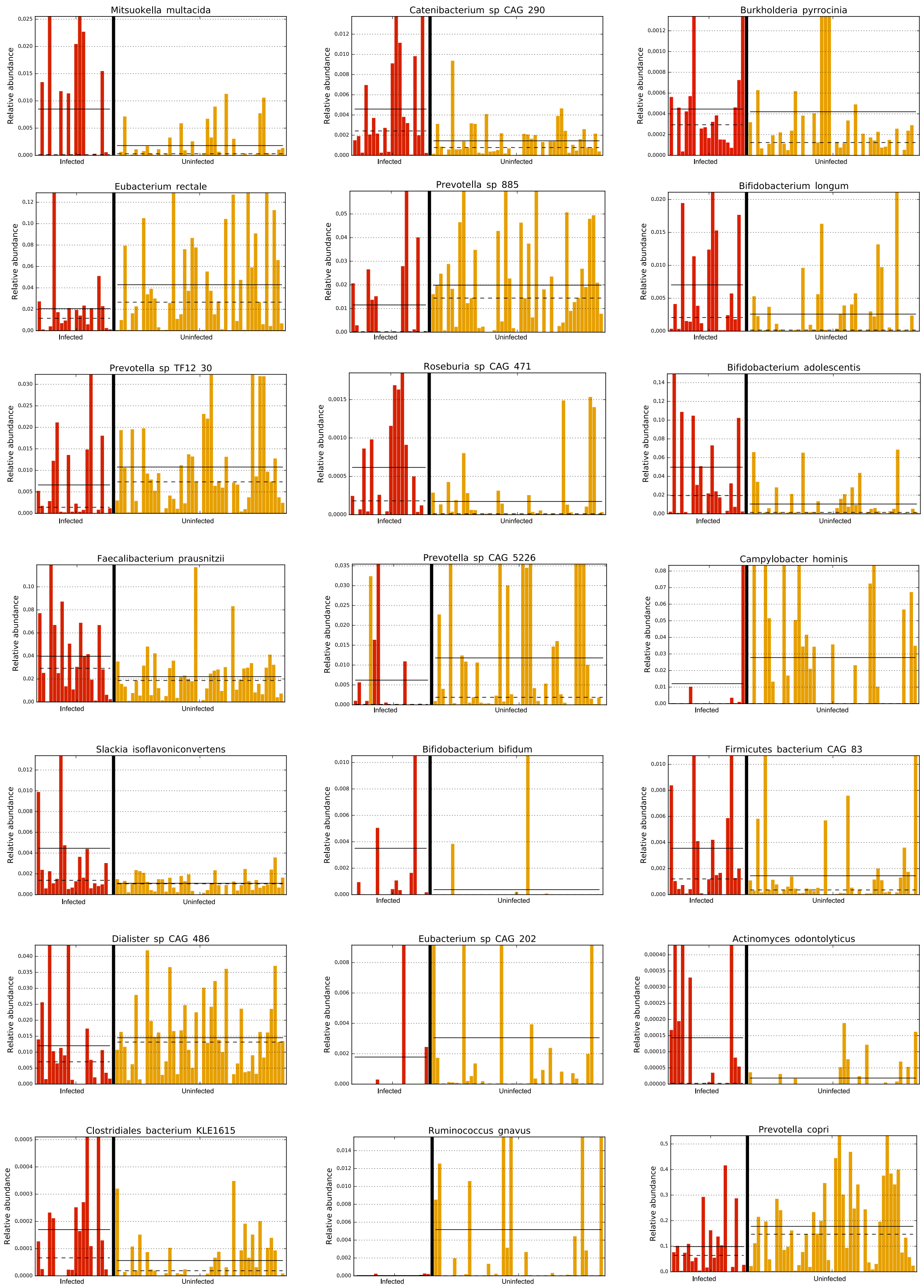


**Fig. S7. Relative abundance of the top 21 most important discriminating species of the gut microbiome of contacts at the time of exposure to *V. cholerae* identified in the Midani 2018 dataset for two classes (Uninfected vs Infected).** The straight line indicates the group means, and the dotted line indicates the group medians.

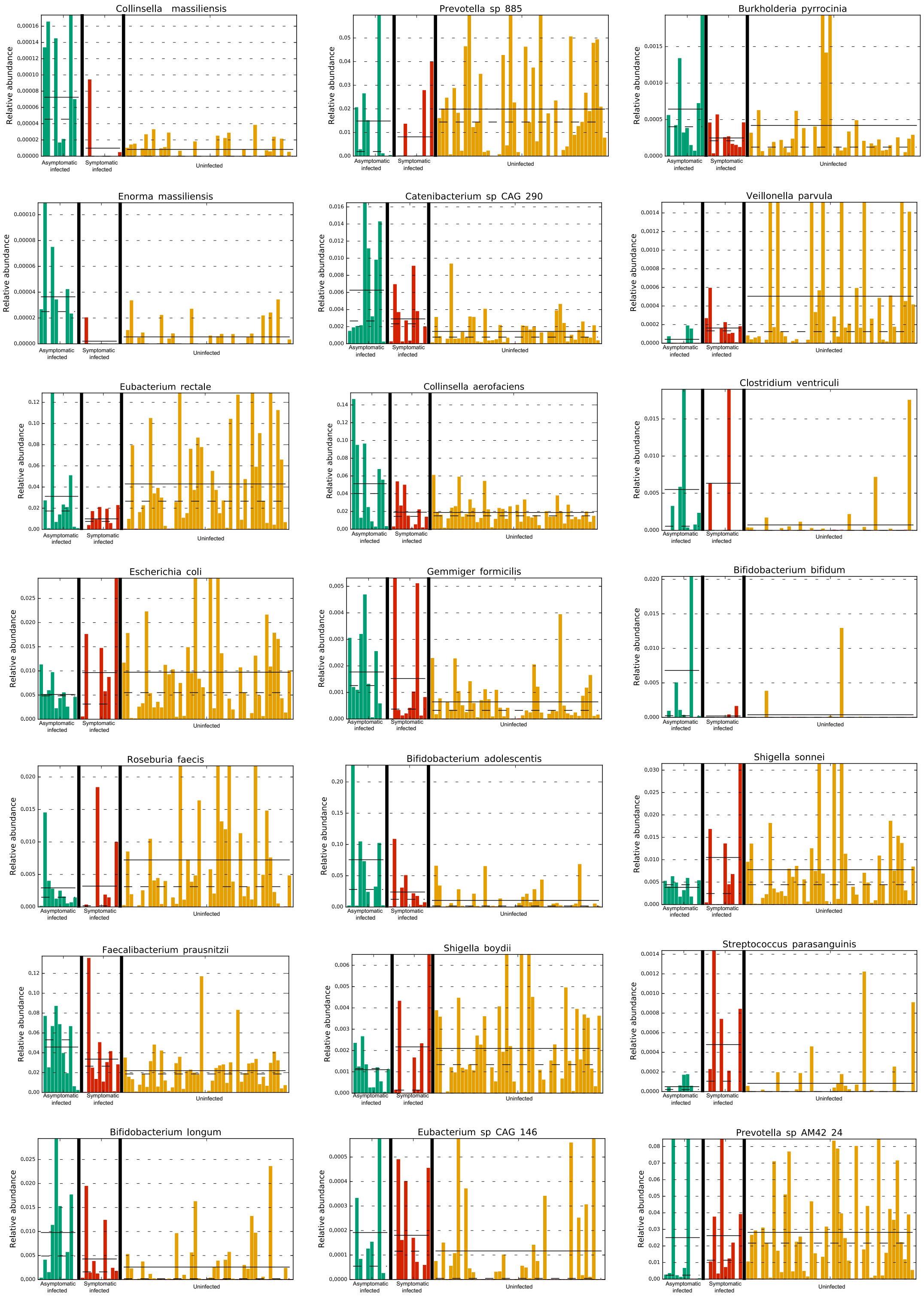


**Fig. S8. Relative abundance of the top 21 most important discriminating species in the gut microbiome of contacts at the time of exposure to *V. cholerae* identified in the Midani 2018 dataset for three classes (Uninfected vs Asymptomatic Infected and Symptomatic Infected).** The straight line indicates the group means, and the dotted line indicates the group medians

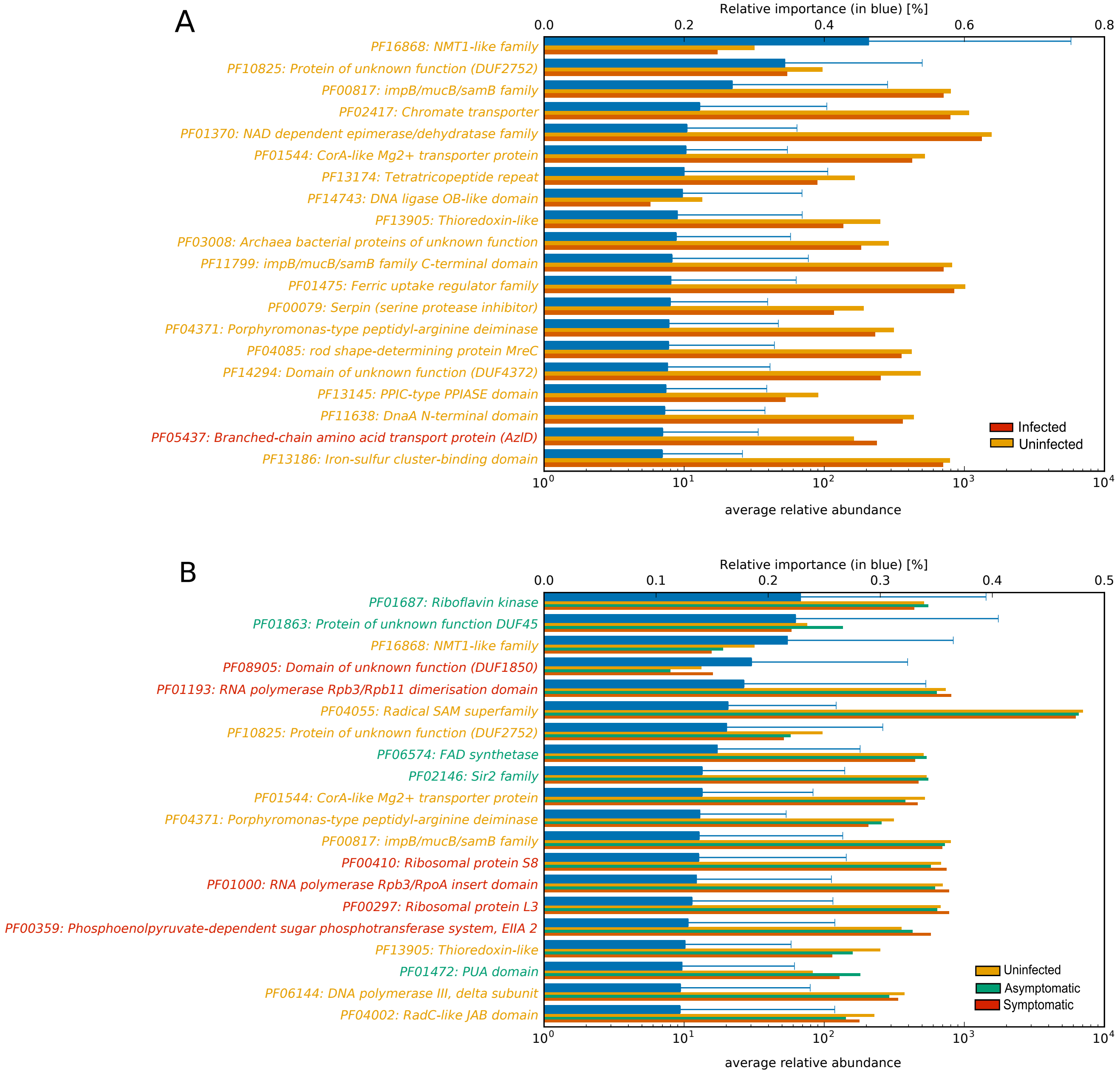


**Figure S9. Most important discriminating gene families of the gut microbiome at the time of exposure to *V. cholerae* identified in the Midani 2018 dataset, classified by clinical outcome.** (A) Genes associated with contacts that became uninfected/infected during follow-up. (B) Genes associated with contacts that became uninfected/asymptomatic/symptomatic during follow-up. For each gene-family reported on the vertical axis, the top bar (in blue) corresponds to the feature relative importance (with standard deviation) and the other bars refer to the average relative abundance (copies per million). The top 25 most important features are shown here; See Table S8 for the full list. Feature relative importance was computed using the mean decrease in impurity strategy, as described in the Methods.

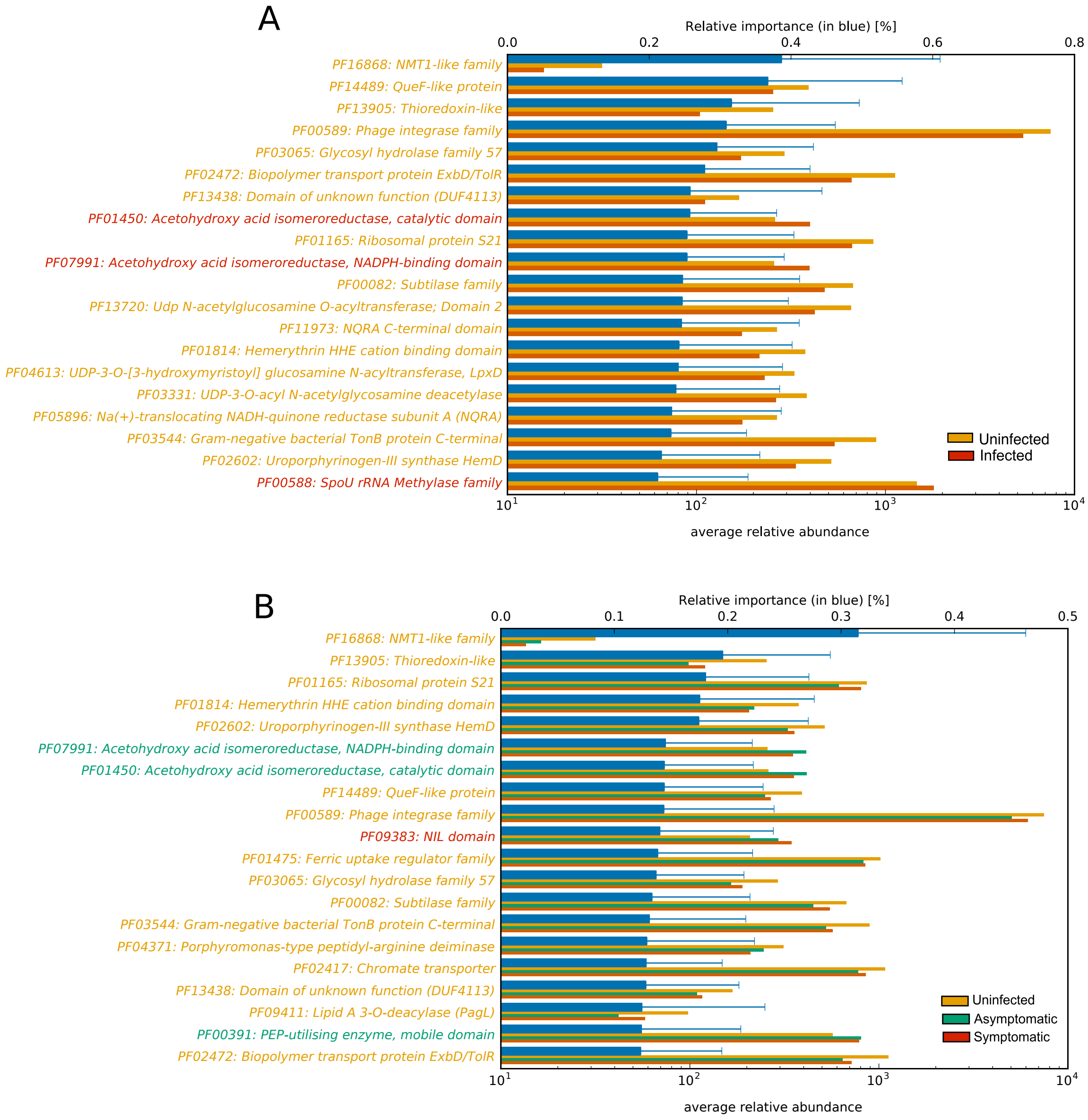


**Fig. S10. Most important discriminating gene-families (grouped by Pfam domain) identified with the random forest algorithm on the Expanded dataset for (A) contacts that became infected or remained uninfected and (B) contacts that remained uninfected, or became infected and were asymptomatic vs symptomatic.** For each gene-families reported on the vertical axis, the top bar (in blue) corresponds to the feature relative importance (with standard deviation) and the other bars refer to the average relative abundance (copies per million). Feature relative importance (blue) is computed using the mean decrease impurity strategy.

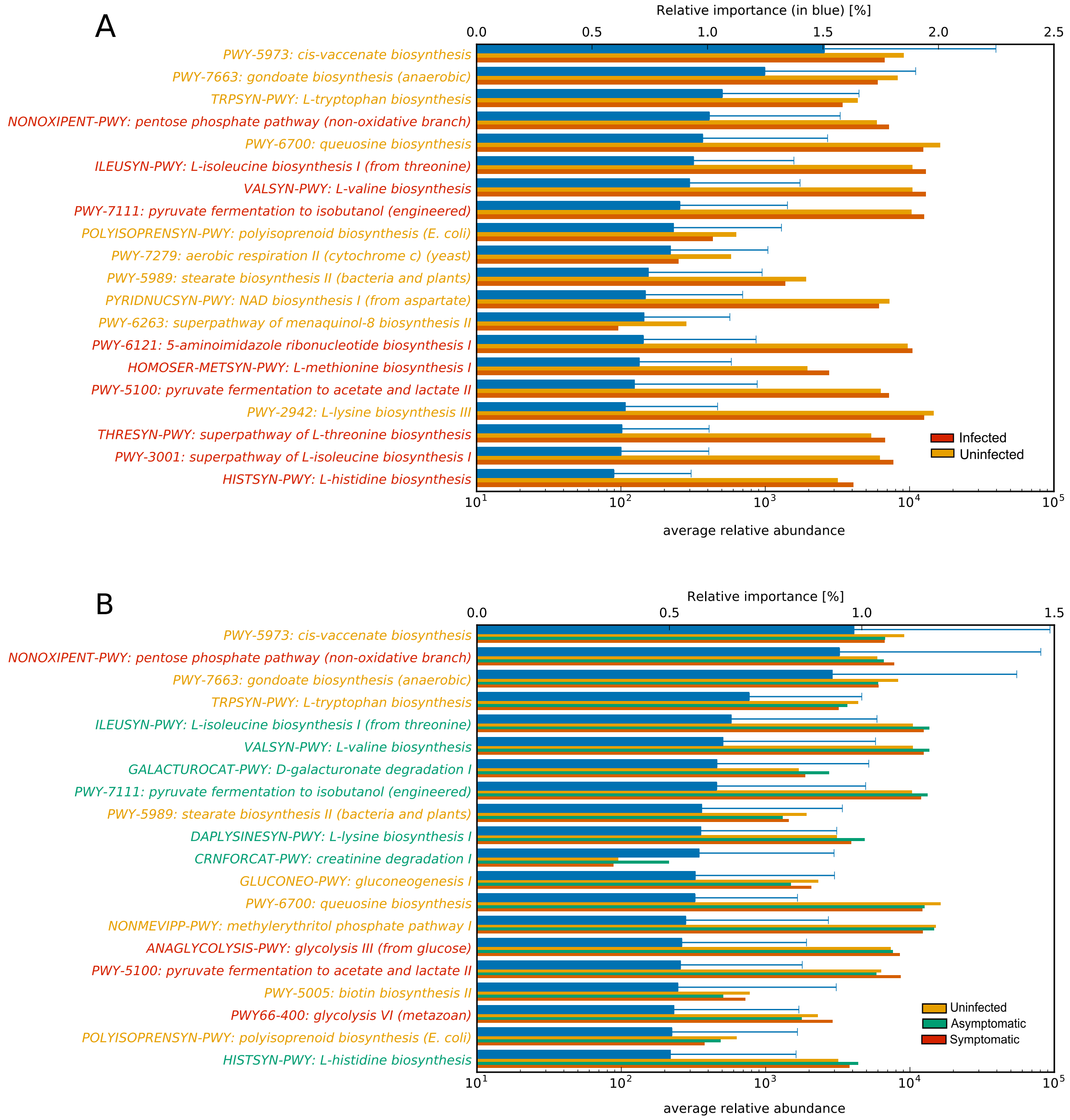


**Fig. S11. Most important discriminating pathways identified with the random forest algorithm on the Midani_2018 dataset for (A) contacts that became infected or remained uninfected and (B) contacts that remained uninfected or became infected and were asymptomatic vs symptomatic.** For each pathway reported on the vertical axis, the top bar (blue) corresponds to the feature relative importance (with standard deviation) and other bars refer to the average relative abundance (copies per million). Feature relative importance (blue) is computed using the mean decrease impurity strategy.

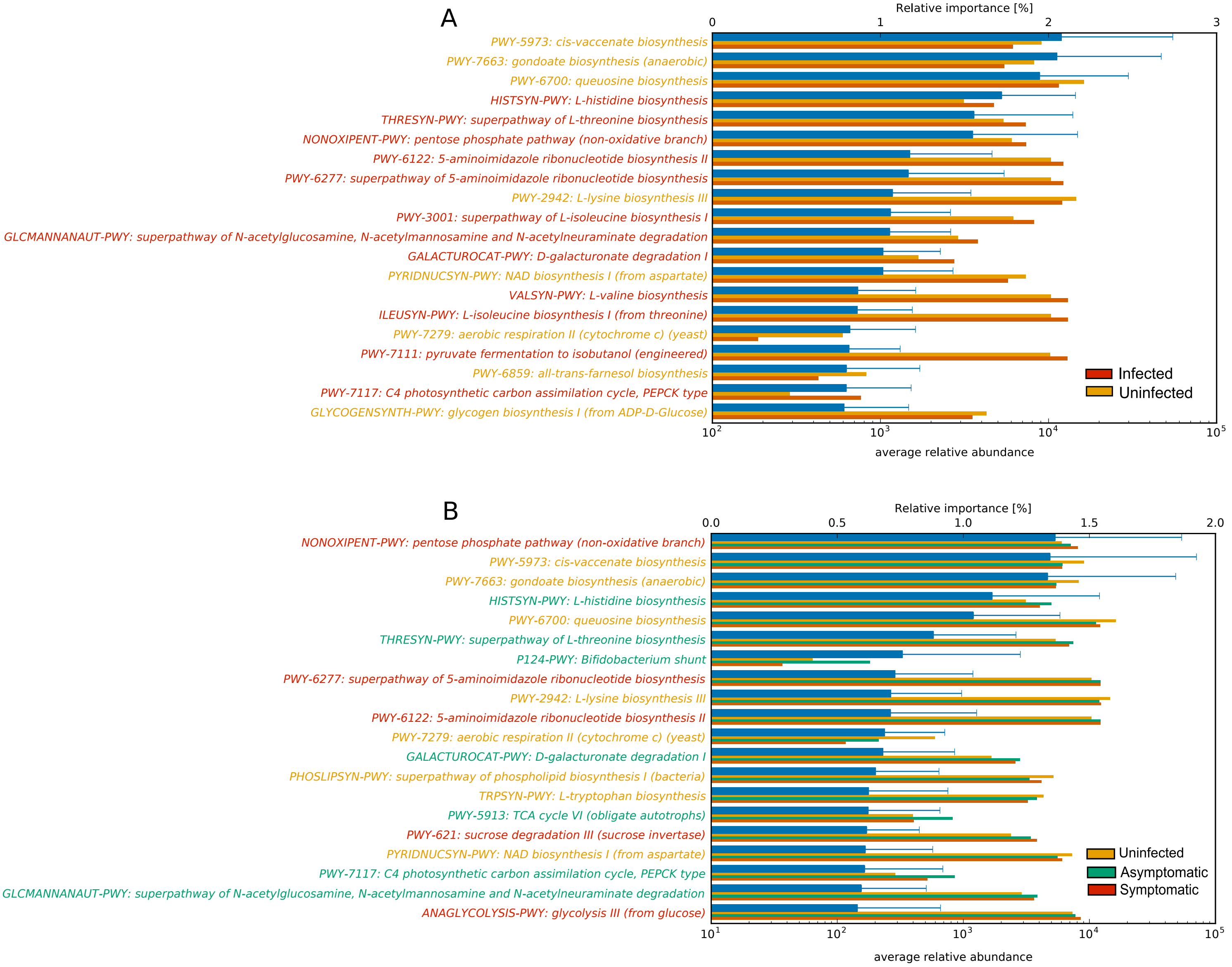


**Fig. S12. Most important discriminating pathways identified with the random forest algorithm on the expanded dataset for (A) contacts that became infected or remained uninfected and (B) contacts that remained uninfected, or became infected and were**

**asymptomatic vs symptomatic.** For each pathway reported on the vertical axis, the top bar (in blue) corresponds to the feature relative importance (with standard deviation) and the other bars refer to the average relative abundance (copies per million). Feature relative importance (blue) is computed using the mean decrease impurity strategy.

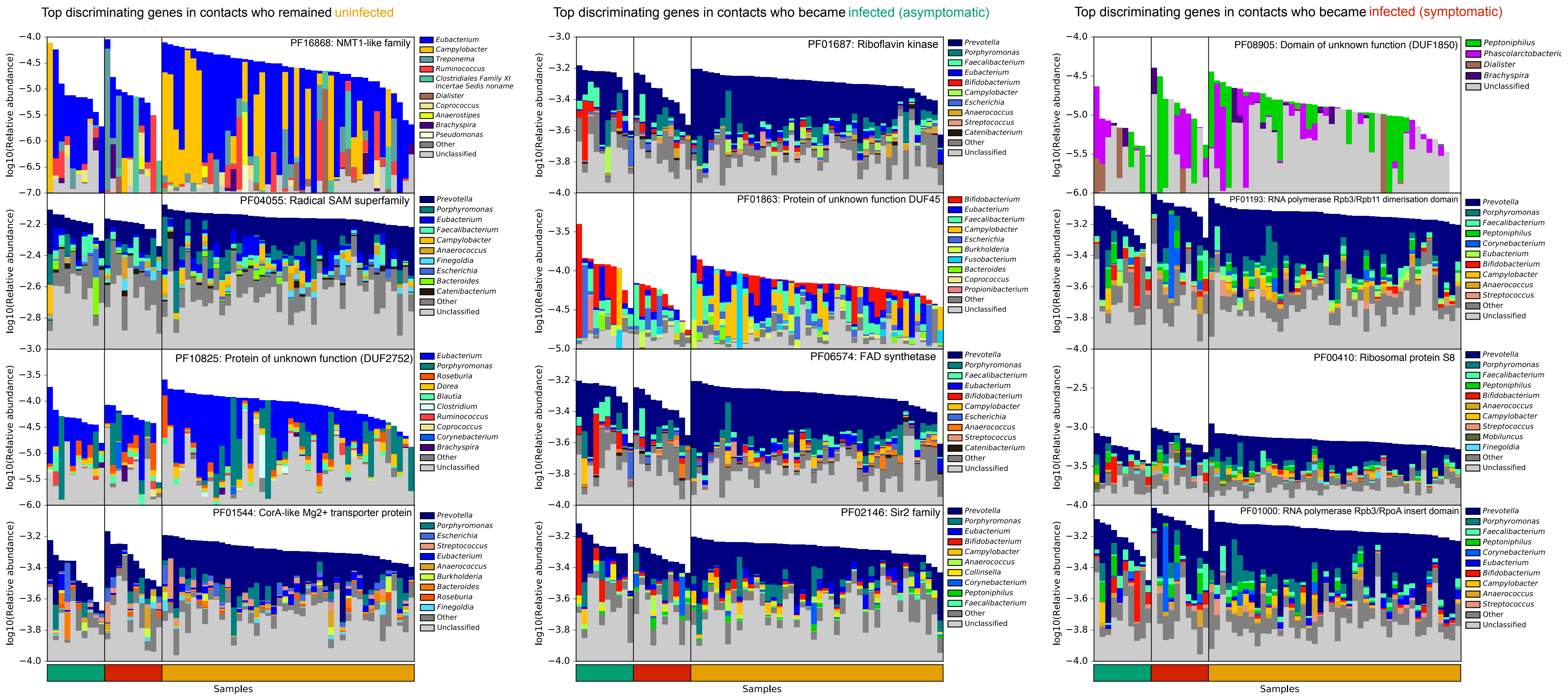


**Fig. S13. Top predictive gene families (Pfam domain grouping) for each class (uninfected, asymptomatic and symptomatic infected contacts), annotated by their taxonomic contributors.** Total bar height reflects $log_{10}$-scaled community abundance. Genera contributions are linearly scaled within total.

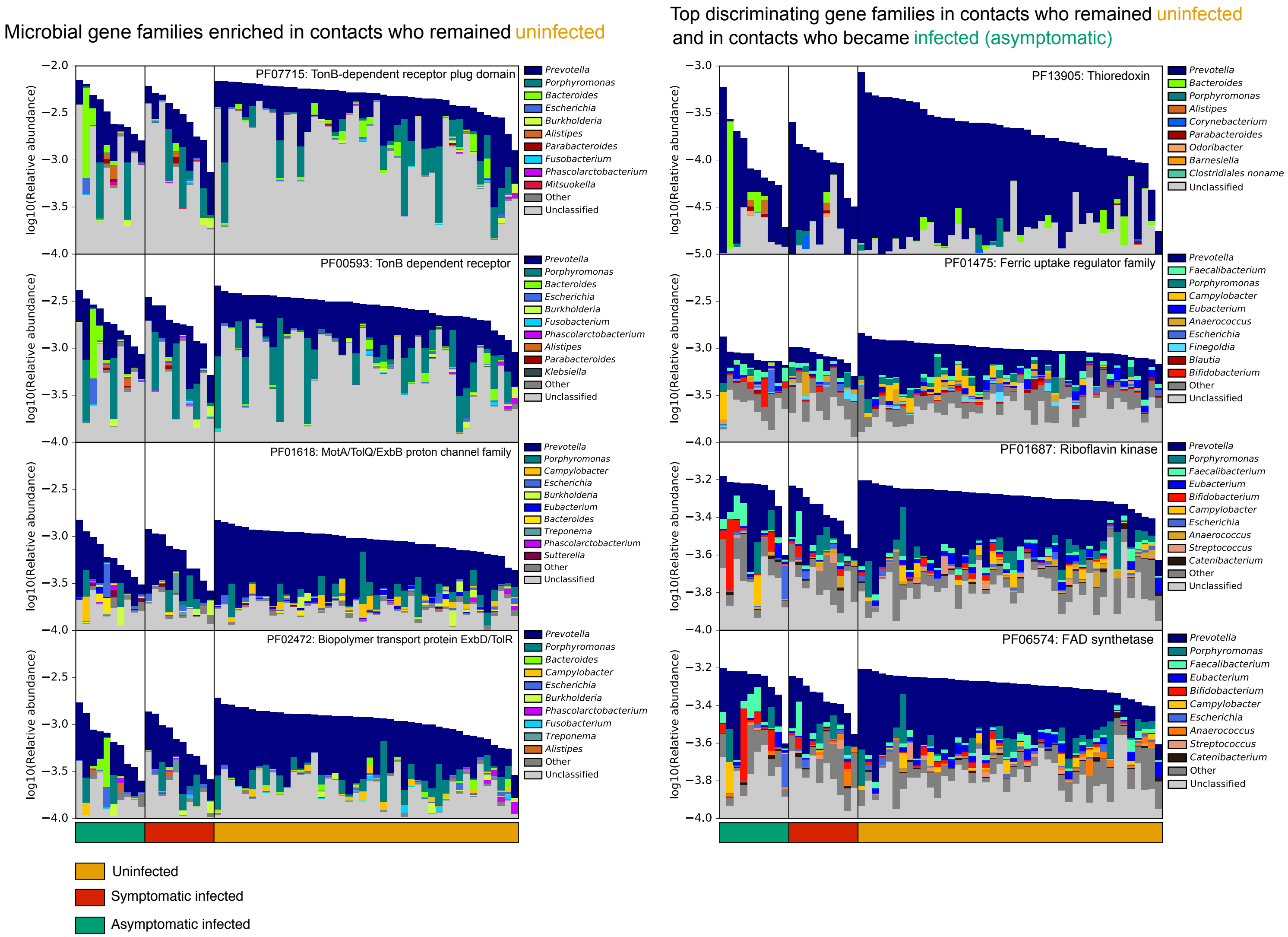


**Fig. S14. Enriched and top predictive gene families (Pfam domain grouping) involved in iron metabolism, annotated by their taxonomic contributors.** Gene families involved with iron metabolism that were differentially abundant in contacts who remained uninfected were identified with LEfSe are represented on the left (shown in Supplementary Fig 2). On the right are the top predictive gene families involved with iron metabolism identified with MetAML. Total bar height reflects $log_{10}$-scaled community abundance. Genera contributions are linearly scaled within total.
